# Endogenous retrovirus rewired the gene regulatory network shared between primordial germ cells and naïve pluripotent cells in hominoids

**DOI:** 10.1101/2021.03.09.434541

**Authors:** Jumpei Ito, Yasunari Seita, Shohei Kojima, Nicholas F. Parrish, Kotaro Sasaki, Kei Sato

**Author notes:** Correspondence (K. Sato), (K. Sasaki). These authors contributed equally (Kotaro Sasaki, Kei Sato).

## Abstract

Although the gene regulatory network controlling germ cell development is critical for gamete integrity, this network has been substantially diversified during mammalian evolution. Here, we show that several hundred loci of LTR5_Hs, a hominoid-specific endogenous retrovirus (ERV), function as enhancers in both human primordial germ cells (PGCs) and naïve pluripotent cells. PGCs and naïve pluripotent cells exhibit a similar transcriptome signature, and the enhancers derived from LTR5_Hs contribute to establishing such similarity. LTR5_Hs appears to be activated by transcription factors critical in both cell types (*KLF4*, *TFAP2C*, *NANOG*, and *CBFA2T2*). Comparative transcriptome analysis between humans and macaques suggested that the expression of many genes in PGCs and naïve pluripotent cells has been upregulated by LTR5_Hs insertions in the hominoid lineage. Together, this study suggests that LTR5_Hs insertions have rewired and finetuned the gene regulatory network shared between PGCs and naïve pluripotent cells during hominoid evolution.

**Teaser:** A hominoid-specific ERV has rewired the gene regulatory network shared between PGCs and naïve pluripotent cells.

## Introduction

Mammalian germ cells are first established as primordial germ cells (PGCs) from pluripotent cells, such as epiblasts, in postimplantation embryos (*1–3*). Aberrations in germ cells lead to immediate infertility, genetic or epigenetic disorders in offspring, and genome integrity impairment. Therefore, the differentiation of germ cells, including PGCs, is strictly controlled by a complex gene regulatory network (*1–3*).

There is an increasing demand to investigate the gene regulatory network using human germ cells. However, it is ethically challenging to routinely access human germ cells, particularly those from humans at early stages of development. Recent studies have established methodologies to derive human germ cells such as PGCs or more differentiated cells from human induced pluripotent stem cells (iPSCs) (*4–7*). These methods have enabled us to characterize the mechanisms of human germ cell development in detail. For example, previous studies using these methods have identified the master regulators of human PGCs, such as *PRDM1*, *SOX17*, *TFAP2C*, and *TFAP2A* (*4, 5, 8, 9*).

The gene regulatory network controlling the development of germ cells such as PGCs is critical for gamete integrity. However, substantial differences exist in this network among mammalian species. For example, various transcription factors (TFs) are differentially expressed between humans and mice (*10*). In particular, *SOX17* is a master regulator of PGC fate specification in humans but not in mice (*4, 5, 8*). Additionally, a substantial number of genes are differentially expressed in PGCs between humans and the crab-eating macaque (*Macaca fascicularis*), Old World monkey (OWM), although the expression patterns of the master regulators of PGCs are conserved between the two species (*11*). These observations suggest that the gene regulatory network controlling germ cell development has been finetuned during mammalian evolution.

Diversification of the gene regulatory network is a molecular basis of evolution and driven by turnover of regulatory sequences such as enhancers (*12, 13*). A substantial proportion of transposable elements (TEs) work as enhancers and play critical roles in the gene regulatory network and its evolution (*14*). Endogenous retroviruses (ERVs) are a class of TEs originating from past retroviral infections. ERVs are particularly rich sources for creation of new enhancers since they contain many regulatory elements in their long terminal repeat (LTR) sequences, which originally function as viral promoters (*15–17*). Notably, since ERV loci belonging to one ERV group share the same set of regulatory elements, numerous inserted ERV loci can coordinately alter the expression patterns of multiple genes (*17–19*). Furthermore, ERVs tend to possess regulatory elements activated in germline niches to proliferate in the germline genome (*17*). Therefore, it is possible that ERVs have been involved in the evolution of the gene regulatory network in germ cells (*20*).

Human PGCs exhibit complex and mixed transcriptome signatures since various gene expression programs are initiated at this stage (*8*). In particular, human PGCs highly express genes associated with naïve pluripotency (*9, 21*). Pluripotency is classified into naïve and primed states, which represent the ground and more-differentiated states, respectively (*22, 23*). Several key TFs, including naïve pluripotency factors (e.g., *NANOG*, *KLF4*, and *TFCP2L1*) and some master regulators of PGCs (e.g., *TFAP2C* and *PRDM1*), are commonly upregulated in human PGCs and naïve pluripotent cells (*4, 5, 8-10, 24-26*). These observations suggest that the core gene regulatory network, which is driven by the key TFs above, might be shared between PGCs and naïve pluripotent cells and play essential roles in establishing cellular identities in these cells. However, this network has not been explored in detail. In particular, the genes and regulatory elements commonly upregulated in PGCs and naïve pluripotent cells have been largely uncharacterized.

In the present study, we investigated the gene regulatory network shared between human PGCs and naïve pluripotent cells in detail. In this process, we found that several hundred loci of LTR5_Hs, the youngest human ERV subfamily expanded in the hominoid lineage (including humans, chimpanzees, gorillas, orangutans, and gibbons, but not OWMs), work as enhancers and play pivotal roles in the gene regulatory network. This study provides evidence suggesting that LTR5_Hs insertions rewired the gene regulatory network shared between PGCs and naïve pluripotent cells during hominoid evolution and possibly accelerated germ cell evolution.

## Results

### Similarity of the gene expression signature of PGCs with that of naïve pluripotent cells

To characterize the transcriptome similarity between PGCs and naïve pluripotent cells, we compared the transcriptome signatures of *in vitro*-derived human PGCs (PGC-like cells; PGCLCs) and naïve embryonic stem cells (ESCs). We analyzed single-cell RNA sequencing (scRNA-Seq) datasets for *in vitro*-derived human male germ cells [Hwang et al. (*7*)] and for naïve and primed ESCs [Messmer et al. (*27*)] (**Fig. 1A**). The Hwang et al. dataset contains information for germ cells that were sequentially differentiated from primed iPSCs: incipient mesoderm-like cells (iMeLCs), PGCLCs, multiplying prospermatogonia-like cells (MLCs), and mitotically quiescent T1 prospermatogonia-like cells (T1LCs), which are formed via transitional cells (TCs) (**Fig. 1A**) (*7*). Dimension reduction analysis suggested that the global transcriptome is highly similar between PGCLCs and naïve ESCs, consistent with previous reports (**Fig. 1A**) (*9, 21*).

**Fig. 1.**
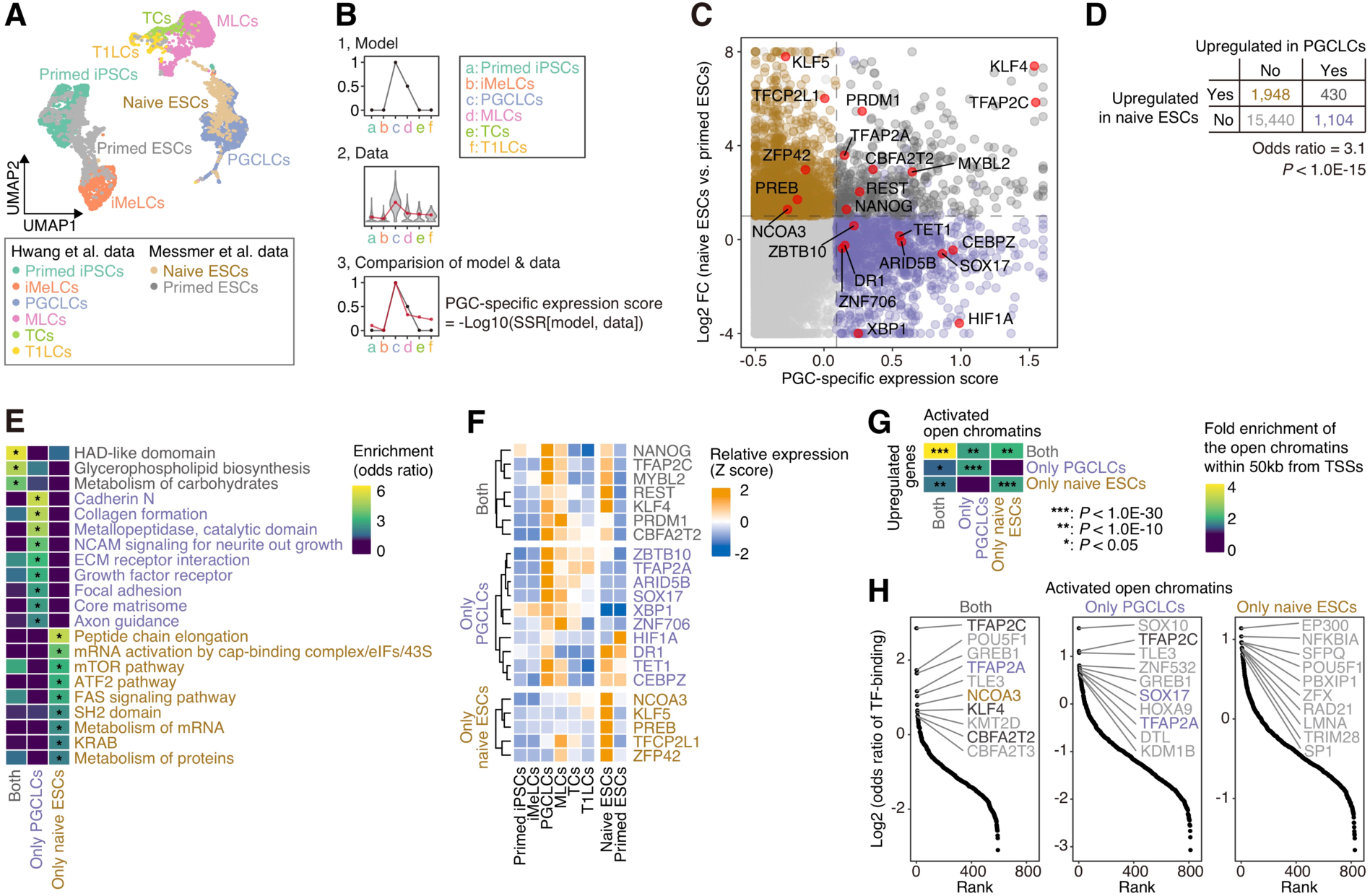
Characterization of the gene expression signature similarity between PGCLCs and naïve ESCs. (A) Dimension reduction analysis of scRNA-Seq data using UMAP (*51*). Data for *in vitro*-derived human male germline development [Hwang et al. (*7*)] and for naïve and primed ESCs [Messmer et al. (*27*)] were integrated and subsequently used. The 3,000 protein-coding genes that were the most differentially expressed among cells were used. (B) Scheme for definition of the PGC-specific expression score. For each gene and TE, the sum of squared residuals (SSR) between the model (panel 1) and the data (i.e., the normalized mean expression value for each cell type; panel 2) was calculated (panel 3). Subsequently, the SSR value was −log10-transformed (see **Methods**). (C) Classification of protein-coding genes according to their expression patterns. The X-axis indicates the PGC-specific expression score, and the Y-axis indicates the log2 FC of the expression score in naïve ESCs vs. primed ESCs. The top 10% of genes with respect to the PGC-specific expression score were regarded as the genes upregulated in PGCLCs. Genes with log2 FC values > 1 and FDR values < 0.05 were regarded as upregulated in naïve ESCs. The genes were classified into four categories: genes upregulated in both cell types (dark gray), genes upregulated only in PGCLCs (purple), genes upregulated only in naïve ESCs (brown) and other genes (light gray). In addition, TFs (except for KZFPs) with elevated expression were annotated (see **Methods**). The plot for KZFPs is shown in **Fig. S2A**. (D) Association of the set of genes upregulated in PGCLCs with that in naïve ESCs. The *P* value was calculated with Fisher’s exact test. (E) GO enrichment analysis results for the three gene categories (genes upregulated in both cell types, genes upregulated only in PGCLCs, and genes upregulated only in naïve ESCs). The gene sets that exhibited significant enrichment (odds ratio >2, FDR < 0.05; denoted by an asterisk) in any of the three gene categories are shown. (F) Expression patterns of the TFs annotated in (**C**). A violin plot visualization is shown in **Fig. S1A**. Although *TFAP2A* was first classified as a gene upregulated in both PGCLCs and naïve ESCs, we reclassified it as a gene upregulated only in PGCLCs since its expression in naïve ESCs was somewhat low (**Figs. 1F, and S1A**). (G) Enrichment of activated open chromatin regions in the vicinities of the upregulated genes. The three categories of open chromatin regions, namely those activated in both cell types, only PGCLCs, and only naïve ESCs compared to primed ESCs, were detected (log2 FC > 1; FDR < 0.05). Subsequently, for the three categories of the open chromatin regions, the degrees of enrichment in the vicinity of (<50 kb from) the genes upregulated in both cell types, upregulated only in PGCLCs and upregulated only in naïve ESCs were calculated using the GREAT scheme (*58*) (see **Method**). The *P* values were calculated with a binomial test. (H) Enrichment of TF-binding events in the open chromatin regions. A publicly available ChIP-Seq dataset provided by the GTRD (*32*) was used. For each TF, the enrichment (odds ratio) of the binding events in the respective categories of open chromatin regions compared to the other open chromatin regions was calculated. Statistical enrichment was calculated using Fisher’s exact test. Of the TFs with FDR values <0.05, the top 10 TFs with respect to the odds ratio are annotated. The upregulated TFs shown in (**C**) and (**F**) are colored.

To further assess the transcriptional similarity between PGCLCs and naïve ESCs, we first focused on the genes upregulated in both cell types. Accordingly, we assigned a PGC-specific expression score for each gene, which represents how the expression pattern is similar to the defined “PGCLC-specific” expression pattern (**Fig. 1B**; see **Methods**). According to this PGC-specific expression score and the log2-transformed fold change (log2 FC) of the expression score between naïve and primed ESCs, we classified the protein-coding genes into four categories: genes upregulated in both cell types, genes upregulated only in PGCLCs, genes upregulated only in naïve ESCs, and other genes) (**Fig. 1C and Table S1)**. As expected, the genes upregulated in PGCLCs substantially overlapped with those upregulated in naïve ESCs, supporting increased transcriptional similarity between these cell types (**Fig. 1D**). Gene Ontology (GO) enrichment analysis showed that the three gene categories were enriched with distinct functional gene sets (**Fig. 1E and Table S2**). Notably, genes related to the “metabolism of carbohydrates” term were enriched among the genes upregulated in both PGCLCs and naïve ESCs (**Fig. 1E**), suggesting that the mode of carbohydrate metabolism is similar between these cell types. Such similarity of a metabolic process between PGCs and naïve pluripotent cells is reminiscent of observations in mice (*28, 29*).

To identify the TFs responsible for the transcriptional similarity between PGCLCs and naïve ESCs, we classified TFs according to their expression patterns (**Figs. 1C and 1F**). Of the key lineage specifiers of PGCLCs (*TFAP2C*, *SOX17*, and *PRDM1*) (*4, 5, 8*), *TFAP2C* and *PRDM1* were upregulated in both PGCLCs and naïve ESCs, while *SOX17* was upregulated only in PGCLCs (**Figs. 1C, 1F, and S1A**). Furthermore, key regulators of pluripotency (*NANOG*, *KLF4*, and *CBFA2T2*) were upregulated in both PGCLCs and naïve ESCs. Moreover, in addition to the native pluripotency-associated TFs (*KLF5*, *TFCP2L1*, and *ZNF42*) (**Figs. 1C, 1F, and S1A**), a substantial number of Krüppel-associated box (KRAB) domain zinc-finger protein (KZFP) family genes were upregulated only in naïve ESCs (**Figs. S2A and S2B**), consistent with the findings of a previous study (*30*). In contrast, the expression of KZFPs was generally low in PGCLCs but gradually upregulated as PGCLCs progressed into later stages of male germ cell development (**Fig. S2C**).

In addition, we analyzed additional transcriptome datasets for PGCLCs [Kojima et al. (*8*) and the newly obtained data] and naïve ESCs [Takashima et al. (*24*) and Theunissen et al. (*31*)] and confirmed that the upregulation of the TFs mentioned above was observed across datasets (**Fig. S1B**).

### Regulatory elements underlying the transcriptional similarity between PGCLCs and naïve ESCs

To identify the regulatory elements underlying the upregulation of genes in both PGCLCs and naïve ESCs, we investigated published datasets from an assay for transposase-accessible chromatin using sequencing (ATAC-Seq) obtained from PGCLCs and naïve/primed ESCs (*21, 30*). We first identified the open chromatin regions (i.e., ATAC-Seq peaks) that were activated in PGCLCs or naïve ESCs compared to primed ESCs and subsequently classified the open chromatin regions into three categories: those activated in both PGCLCs and naïve ESCs, those activated only in PGCLCs, and those activated only in naïve ESCs. Finally, we examined the enrichment of the different categories of open chromatin regions in the vicinity of (<50 kb from) the genes upregulated in both PGCLCs and naïve ESCs (**Fig. 1G**). The open chromatin regions activated in both cell types were clearly enriched near the genes upregulated in both cell types, suggesting that the regulatory sequences activated in both cell types are particularly important for controlling the upregulated genes common to these cell types (**Fig. 1G**).

To identify the TFs critical for controlling the regulatory elements identified above, we analyzed a publicly available chromatin immunoprecipitation sequencing (ChIP-Seq) dataset for 1,308 types of TFs provided by the Gene Transcription Regulation Database (GTRD) (*32*). For the various TFs, we computed the enrichment of the binding events in each category of open chromatin regions compared to the other identified open chromatin regions (**Fig. 1H**). The open chromatin regions activated in both PGCLCs and naïve ESCs were preferentially bound by TFs that were upregulated in both cell types (*TFAP2C*, *KLF4*, and *CBFA2T2*) or in one of these cell types (*TFAP2A* for PGCLCs and *NCOA3* for naïve ESCs). This result supports the importance of these TFs in regulating the genes upregulated in both cell types (**Fig. 1H**).

### TEs that are commonly upregulated in PGCLCs and naïve ESCs

To identify the TEs that are activated as enhancers during human male germline developmental process, including PGCs, we analyzed the expression dynamics of TEs using the Hwang et al. scRNA-Seq dataset (**Fig. 2**) (*7*). We first used transcriptome data instead of epigenomic data since the transcriptional activity of TEs is known to reflect enhancer activity, similar to the case for enhancer RNAs (*33*). Pseudotime analysis (*34*) showed that the expression of TEs dynamically changed during *in vitro*-derived male germline development (**Figs. 2A and B**). As described previously (*7*), the expression of most TEs (long interspersed nuclear elements [LINEs], short interspersed nuclear elements [SINEs], and SINE-VNTR-Alu [SVA] and DNA transposons) was gradually upregulated with the progression of development, presumably reflecting the gradual DNA demethylation that occurred during this process (**Fig. 2B**) (*10, 25*). On the other hand, the expression of the various ERV subfamilies, including HERVH, LTR7, and LTR12C, was stage-specific (**Fig. 2B**) (*7*). In particular, the expression of some ERV subfamilies, such as HERVK, LTR5_Hs and HERVIP10FH, was specifically upregulated in PGCLCs and subsequently downregulated in cells at later stages (i.e., MLCs, TCs, and T1LCs) (**Figs. 2B and 2C**). Notably, HERVK/LTR5_Hs (LTR5_Hs is a type of the LTR sequence of HERVK) was one of the top-ranked TEs with respect to the PGC-specific score (**Fig. 2D, X-axis**). On the other hand, SVA transposons, a group of chimeric TEs originating partially from HERVK/LTR5_Hs (*35*), did not exhibit such a PGC-specific expression pattern (**Figs. 2B, 2D, and S3**).

**Fig. 2.**
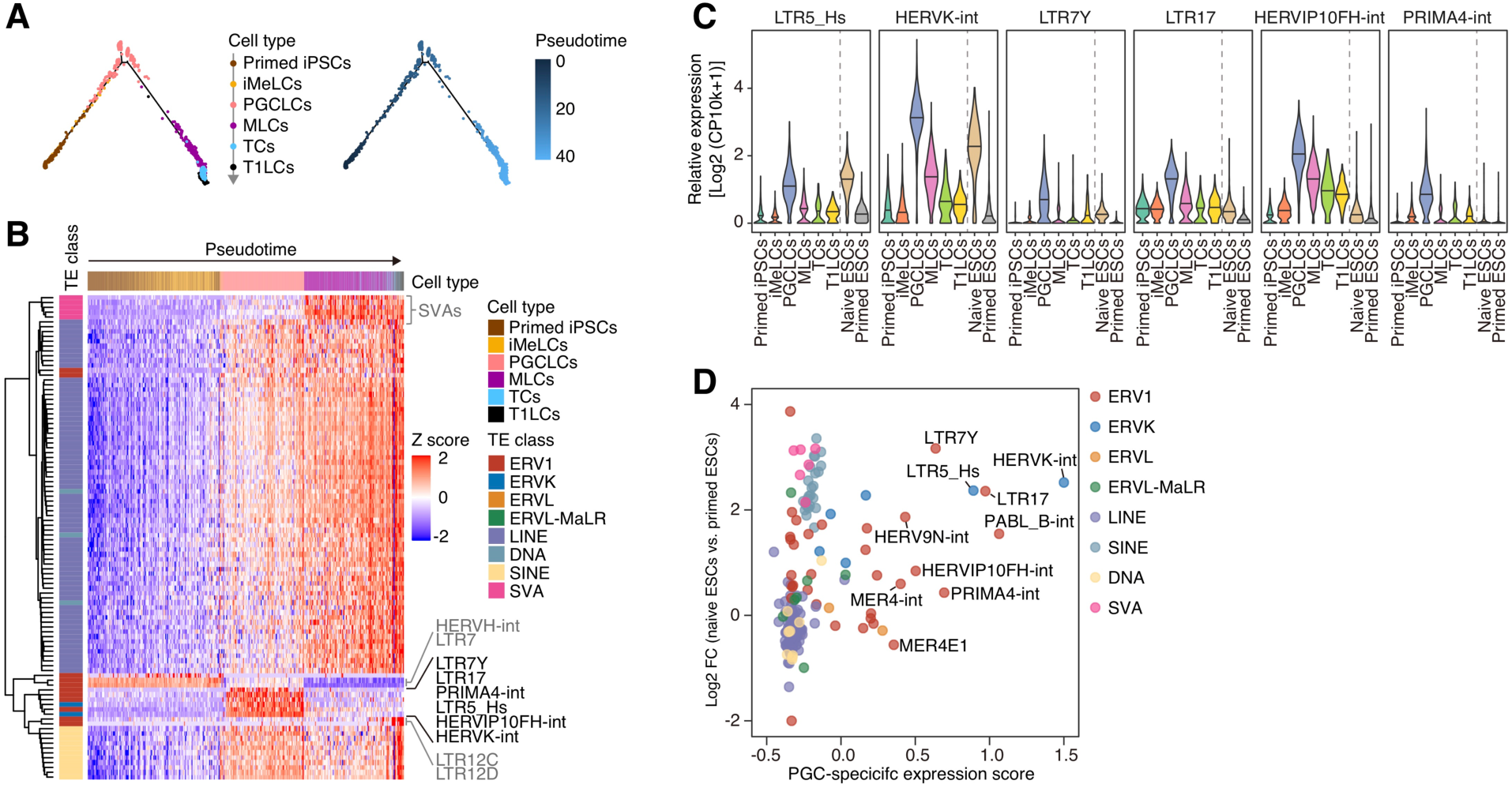
Specific expression of HERVK/LTR5_Hs in PGCLCs and naïve ESCs. (A) Pseudotime analysis (*34*) of scRNA-Seq data for *in vitro*-derived human male germline development [Hwang et al. (*7*)]. The 1,000 protein-coding genes that were the most differentially expressed throughout the development process were used. (B) Expression dynamics of TE subfamilies throughout male germline development. The cells are ordered according to the pseudotime shown in (**A**). The 100 TEs that were most differentially expressed among cell types are shown. (C) ERV subfamilies that were specifically expressed in PGCLCs [annotated in (**B**) in black]. In addition to the data for male germline development, data for naïve and primed ESCs [Messmer et al. (*27*)] are shown. (D) Identification of the TE subfamilies that were specifically upregulated in both PGCLCs and naïve ESCs. The X-axis indicates the PGC-specific expression score (defined in Fig. 1B). The Y-axis indicates the log2 FC of the expression score between naïve ESCs vs. primed ESCs. The names of the top 10% TEs with respect to the PGC-specific expression score are annotated.

Previous studies have shown that HERVK/LTR5_Hs is highly activated in naïve pluripotent cells, such as naïve ESCs and cells in the inner cell masses of blastocysts (**Fig. 2C**) (*30, 31, 36, 37*). Indeed, our data showed that HERVK/LTR5_Hs was one of the top-ranked TEs upregulated in both PGCLCs and naïve ESCs (**Fig. 2D**). Together, these results raise the possibility that LTR5_Hs may serve as enhancers shared between PGCs and naïve pluripotent cells and contribute to establishing the transcriptional similarity between these two cell types.

### Increased enhancer activity of LTR5_Hs in PGCLCs and naïve ESCs

To evaluate the enhancer potential of LTR5_Hs in PGCLCs and naïve ESCs, we investigated the chromatin accessibility and histone modification status of LTR5_Hs in these two cell types using ATAC-Seq and ChIP-Seq data targeting an active histone mark (i.e., H3K27ac), respectively (**Fig. 3**). We examined the statistical enrichment of the two types of epigenetic signals on TEs in the various subfamilies (**Figs. 3A and 3B**). In terms of both chromatin accessibility and active histone marks, LTR5_Hs was the top-ranked TE that was epigenetically activated in both PGCLCs and naïve ESCs. We next examined whether the chromatin accessibility of LTR5_Hs was greater in PGCLCs and naïve ESCs than in primed ESCs (**Fig. 3C**). The open chromatin regions overlapping with LTR5_Hs tended to be activated in PGCLCs (**Fig. 3C, upper panel**) and naïve ESCs (**Fig. 3C, right panel**) compared to primed ESCs.

**Fig. 3.**
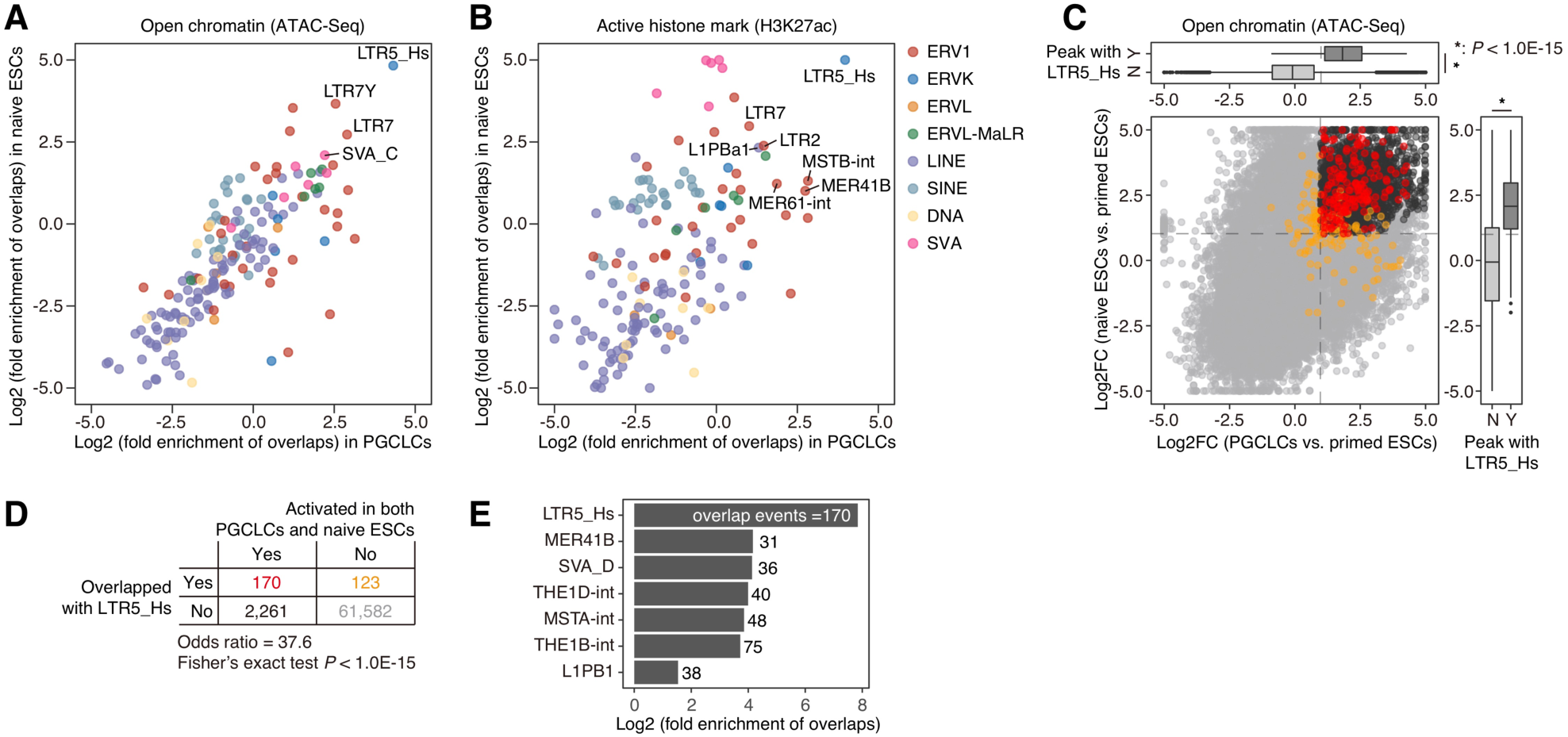
Potential enhancer activity of LTR5_Hs in PGCLCs and naïve ESCs. (A and B) Fold enrichment of the genomic overlap between TE loci and the peaks of ATAC-Seq (**A**) and ChIP-Seq targeting an active histone mark, H3K27ac (**B**). The fold enrichment value compared to the random expectation was calculated by the genomic permutation test. The X-axis and Y-axis indicate the log2-transformed fold enrichment values in PGCLCs and naïve ESCs, respectively. The ATAC-Seq and ChIP-Seq data originated from Pontis et al. (*30*) and Chen et al. (*21*). (C) Upregulation of the chromatin accessibility of LTR5_Hs loci in PGCLCs and naïve ESCs compared to primed ESCs. For each ATAC-Seq peak (i.e., open chromatin region), the log2 FC scores of the chromatin accessibility in PGCLCs vs. primed ESCs (the X-axis) and naïve ESCs vs. primed ESCs (the Y-axis) are shown. In the main panel, the peaks overlapping with LTR5_Hs are colored red or orange. The peaks are colored red or black if they were upregulated in both PGCLCs and naïve ESCs (log2 FC > 1; FDR < 0.05). The color scheme is summarized in (**D**). In the upper and right panels, the marginal distributions respectively for the X- and Y-axes are shown (Y [Yes], overlapped with LTR5_Hs; N [No], not overlapped). An asterisk denotes *P* < 1.0E-15 in the two-tailed Wilcoxon rank sum test. (D) The enrichment of LTR5_Hs in the ATAC-Seq peaks upregulated in both PGCLCs and naïve ESCs compared to primed ESCs. The *P* value was calculated with Fisher’s exact test. (E) The enrichment of the various TE subfamilies in the ATAC-Seq peaks was upregulated in both PGCLCs and naïve ESCs. The fold enrichment value compared to the random expectation and the statistical significance were computed with the genomic permutation test. The number of overlap events is shown on each bar. The results for TEs with significant enrichment (FDR < 0.05; log2 fold enrichment > 1; overlap events > 20) are shown.

Furthermore, LTR5_Hs was highly enriched in the open chromatin regions that were significantly activated in both PGCLCs and naïve ESCs (**Fig. 3C, main panel and Fig. 3D**). Indeed, LTR5_Hs exhibited the strongest enrichment in these commonly activated open chromatin regions among all TEs (**Fig. 3E**). Together, our findings demonstrate that LTR5_Hs serves as an enhancer shared between PGCLCs and naïve ESCs.

### Potential regulators of LTR5_Hs in PGCs and naïve pluripotent cells

We next surveyed the TFs that bind to LTR5_Hs and control its activity in PGCLCs and naïve ESCs (**Fig. 4 and Table S3**). We analyzed the publicly available ChIP-Seq dataset for 1,308 types of TFs and identified TFs that preferentially bound to LTR5_Hs. Of these TFs, we extracted TFs that were expressed specifically in PGCLCs and naïve ESCs (**Figs. 4A and 4B**). Of the TFs that preferentially bound to LTR5_Hs, *NANOG*, *TFAP2C*, *KLF4*, and *CBFA2T2* were upregulated in both PGCLCs and naïve ESCs (**Figs. 1C, 4C, and 4D**). Furthermore, *SOX17* and *TFAP2A* were specifically upregulated in PGCLCs, while *KLF5* was upregulated in naïve ESCs (**Figs. 1C, 4C, and 4D**). Notably, these TFs are known to play central roles in gene regulation in PGCs (i.e., *SOX17* and *TFAP2A*) (*5, 9*), naïve pluripotent cells (i.e., *KLF5*) (*38, 39*) or both cell types (i.e., *NANOG*, *TFAP2C*, *KLF4*, and *CBFA2T2*) (**Fig. 1H**) (*4, 5, 8-10, 24, 26, 40*). Together, our data suggest that the enhancer activity of LTR5_Hs in PGCs and naïve pluripotent cells appears to be controlled by key TFs in these cell types.

**Fig. 4.**
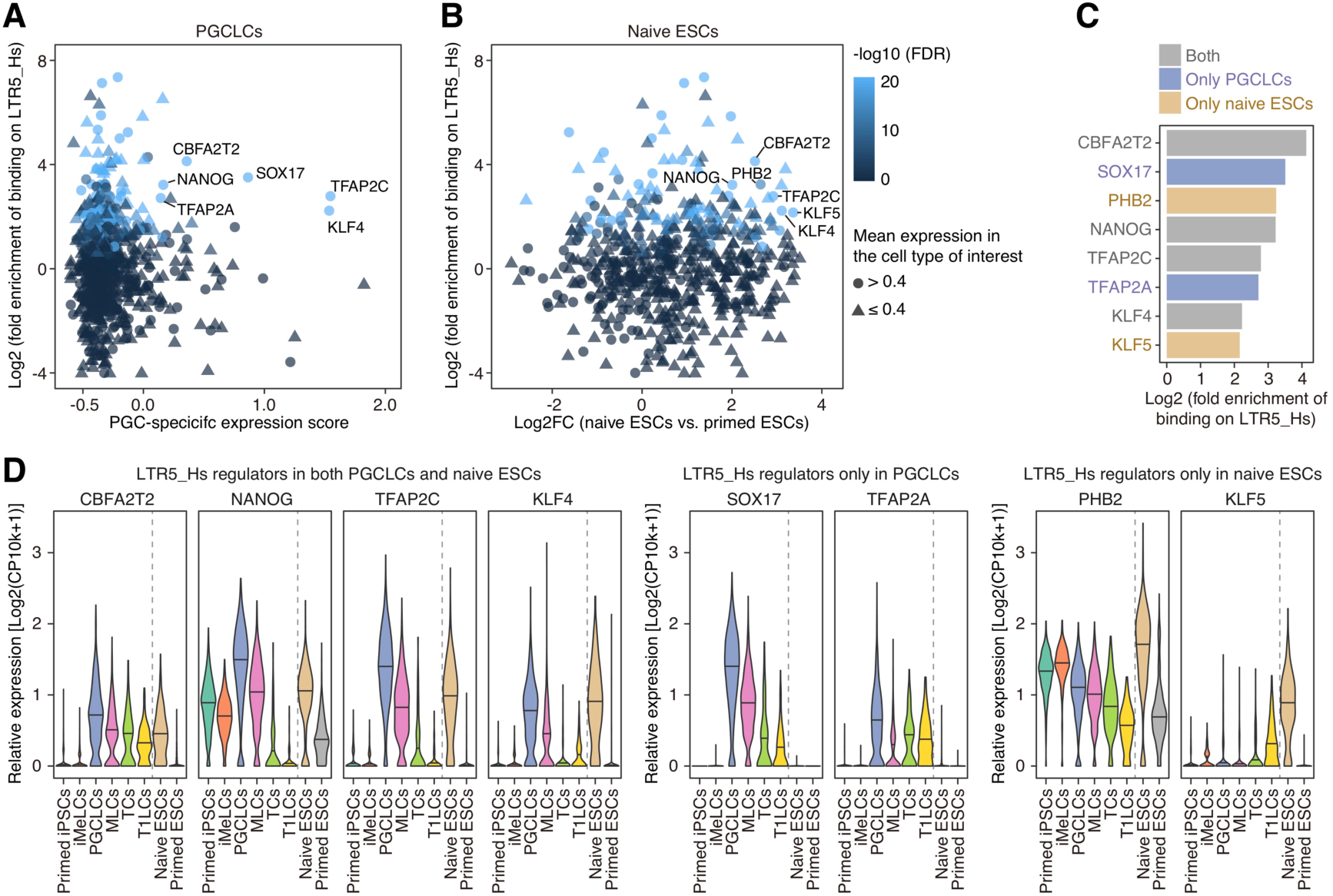
Identification of the potential regulators of LTR5_Hs in PGCLCs and naïve ESCs. (A and B) Identification of the TFs that bind to LTR5_Hs and are upregulated in PGCLCs (**A**) and naïve ESCs (**B**). For each TF, the statistical enrichment of the binding events on LTR5_Hs was calculated based on the publicly available ChIP-Seq dataset provided by the GTRD (*32*). The Y-axis indicates the log2-transformed fold enrichment of the TF-binding events compared to the random expectation. The X-axis indicates the PGC-specific expression score (**A**) or the log2 FC of the expression score in naïve ESCs vs. primed ESCs (**B**). The symbols are colored according to the statistical significance of the TF-binding enrichment calculated by the genome permutation test. The symbol shape represents the mean expression level in PGCLCs (**A**) and naïve ESCs (**B**). The potential regulators of LTR5_Hs are annotated. The potential regulators were defined as the TFs satisfying the following criteria: (i) TFs that exhibited significant binding enrichment on LTR5_Hs (log2 fold enrichment > 2; FDR < 0.05; binding events > 20); (ii) for regulators in PGCLCs, TFs that were specifically upregulated in PGCLCs (the top 10% TFs with respect to the PGC-specific expression score; mean relative expression (log2[CP10k+1] > 0.4 in PGCLCs); and (iii) for regulators in naïve ESCs, TFs that were specifically upregulated in naïve ESCs (log2 FC > 2; FDR < 0.05; mean relative expression > 0.4 in naïve ESCs). (C) Classification of the potential LTR5_Hs regulators. The X-axis indicates the log2-transformed fold enrichment of the TF-binding events. (D) Expression patterns of TFs identified as potential LTR5_Hs regulators.

### Expression patterns of the genes adjacent to LTR5_Hs in PGCLCs and ESCs

To elucidate the roles of LTR5_Hs in gene regulation in PGCs and naïve pluripotent cells, we investigated the expression patterns of the genes adjacent to (<50 kb from) the LTR5_Hs loci with transcriptomic or epigenetic activity (**Fig. 5 and Table S4**). The genes adjacent to LTR5_Hs tended to be specifically upregulated in both PGCLCs (**Fig. 5A, upper panel**) and naïve ESCs (**Fig. 5A, right panel**). Notably, the genes adjacent to LTR5_Hs were strikingly enriched with genes upregulated in both PGCLCs and naïve ESCs (**Fig. 5A, main panel and Fig. 5B**). Indeed, of the genes commonly upregulated in PGCLCs and naïve ESCs, approximately 25% (107/430) were located in the vicinity of LTR5_Hs (**Fig. 5B**). These results suggest that LTR5_Hs upregulates adjacent genes as an enhancer in these cell types. GO enrichment analysis showed that genes associated with the “glucose metabolism” and “glycogen breakdown” terms were particularly enriched among the genes adjacent to LTR5_Hs and upregulated in both cell types (**Fig. 5C and Table S5**). These are child terms of the “metabolism of carbohydrates” term, which was significantly enriched for the genes upregulated in both PGCLCs and naïve ESCs (**Fig. 1E**). Furthermore, the glucose metabolism-related genes (i.e., *AGL*, *ENO2*, *PFKL*, *PHKA1*, and *PYGB*) were highly expressed in both PGCLCs and naïve ESCs (**Fig. 5D**). The genes play central roles in energy generation via glycolysis (*PHKA1* and *PYGB*) and glycogenolysis (*AGL*, *ENO2*, and *PFKL*) (**Fig. S4**). These results suggest that enhancers derived from LTR5_Hs play a critical role in the regulation of glucose metabolism in both PGCs and naïve pluripotent cells (see **Discussion**).

**Fig. 5.**
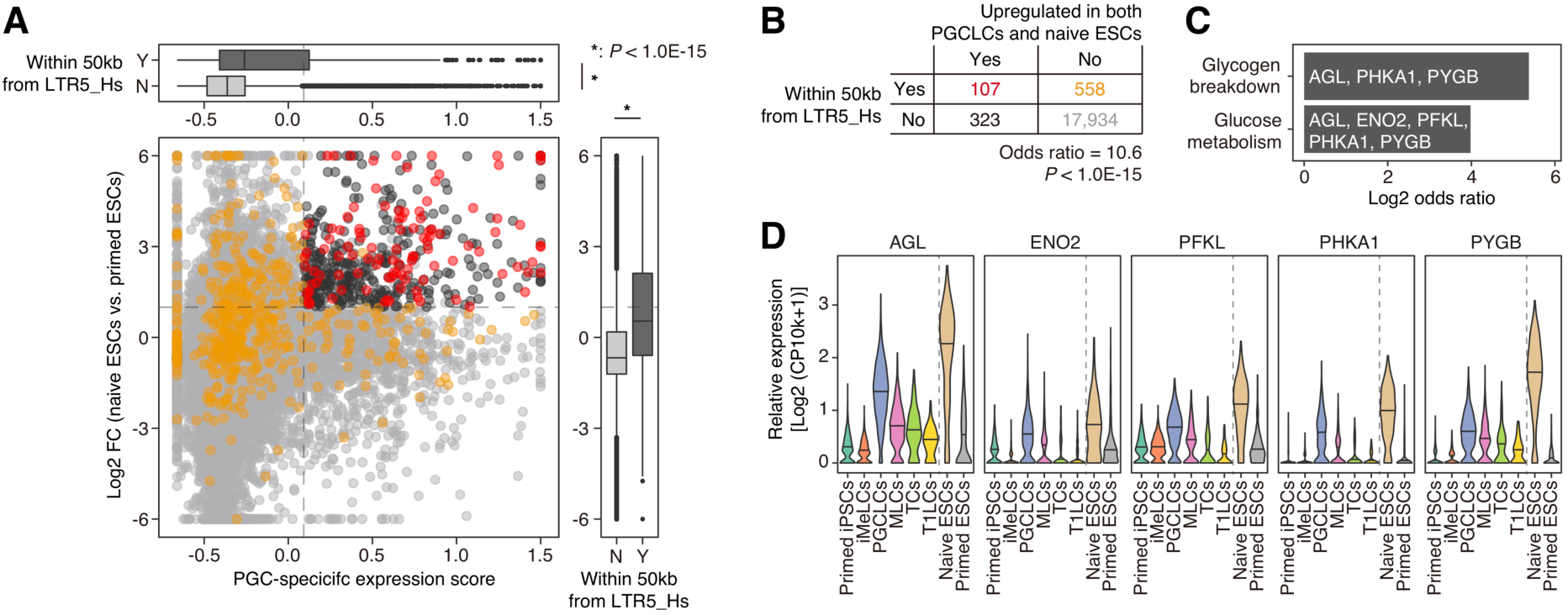
Expression patterns of the genes adjacent to LTR5_Hs in PGCLCs and naïve ESCs. (A) Association of the expression patterns of genes and their distance from LTR5_Hs in the genome. The X-axis indicates the PGC-specific expression score, and the Y-axis indicates the log2 FC of the expression score in naïve ESCs vs. primed ESCs. Genes were stratified according to whether they were present within 50 kb of LTR5_Hs with epigenetic or transcriptomic signals. In the main panel, the genes in the vicinity of LTR5_Hs are colored red or orange. The genes are colored red or black if they were upregulated in both PGCLCs (the top 10% of genes with respect to the PGC-specific expression score) and naïve ESCs (log2 FC > 1; FDR < 0.05). The color scheme is summarized in (**B**). In the top and right panels, the marginal distributions respectively for the X- and Y-axes are shown (Y [Yes], adjacent to LTR5_Hs; N [No], not adjacent). An asterisk denotes *P* < 1.0E-15 in the two-tailed Wilcoxon rank sum test. (B) Enrichment of the genes adjacent to LTR5_Hs among the genes upregulated in both PGCLCs and naïve ESCs. The *P* value was calculated with Fisher’s exact test. (C) Results of GO enrichment analysis. The gene sets with significant enrichment (FDR < 0.05) are shown. The names of the hit genes are shown on each bar. (D) Expression patterns of the genes present in the vicinity of LTR5_Hs and related to glucose metabolism.

### Gene expression alterations driven by LTR5_Hs during primate evolution

LTR5_Hs proliferated in hominoid genomes after the divergence of hominoids and OWMs (*17*). To elucidate the alterations in gene expression driven by LTR5_Hs insertions, we performed comparative transcriptome analysis between humans and an OWM, the crab-eating macaque, focusing on PGCs and naïve pluripotent cells (**Fig. 6**). Similar to the findings in **Fig. 5A**, the results revealed that genes adjacent to LTR5_Hs in the human genome tended to be upregulated commonly in PGCLCs and naïve ESCs compared to primed ESCs (**Figs. 6A and 6C**). On the other hand, the macaque orthologs of the human genes adjacent to LTR5_Hs did not show such a clear tendency (**Figs. 6B and 6D**). Furthermore, the genes upregulated in both PGCs/PGCLCs and naïve pluripotent cells did not highly overlap between humans and macaques (12%, 61/512 in humans), although the upregulation of key TFs, such as *KLF4*, *NANOG*, *TFAP2C*, *PRDM1*, and *CBFA2T2*, was conserved between the two species (**Fig. 6E and Table S6**). Moreover, of the genes that were upregulated in both PGCs/PGCLCs and naïve pluripotent cells only in humans, approximately 21% (95/451) were in the vicinity of LTR5_Hs (**Fig. 6E**). We hereafter refer to these 95 genes as the genes that are likely to be regulated by LTR5_Hs (**Fig. 6E**). Taken together, these results suggest that LTR5_Hs insertions have altered the expression patterns of their adjacent genes to the PGC- and naïve-specific patterns in the hominoid lineage.

**Fig. 6.**
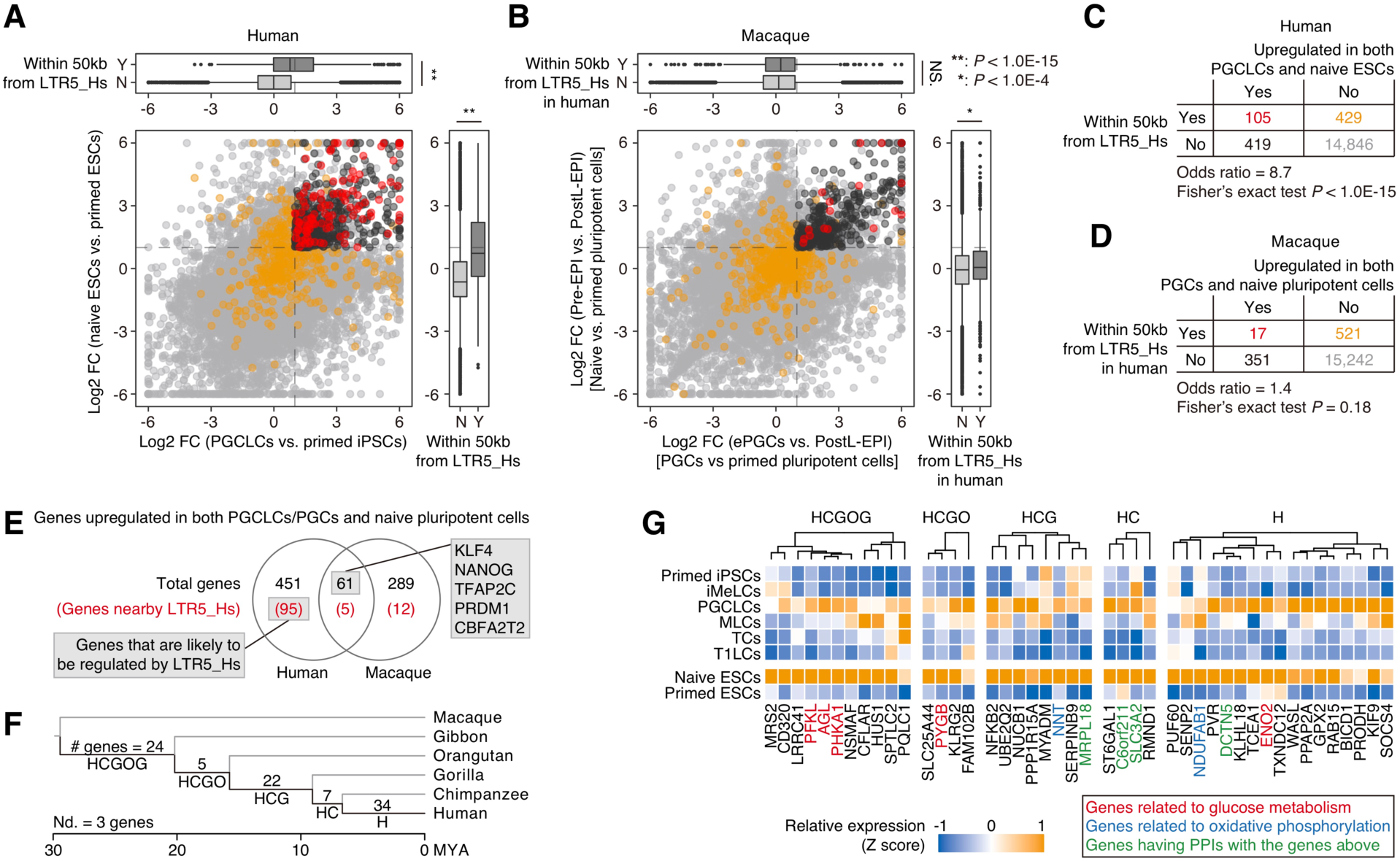
Comparative transcriptome analysis between humans and crab-eating macaques. (A and B) Comparative analysis of the gene expression patterns in PGCLCs/PGCs and naïve pluripotent cells between humans (**A**) and crab-eating macaques (**B**). (**A**) is similar to Fig. 5A, but the X-axis indicates the log2 FC of the expression score in PGCLCs vs. primed iPSCs. In (**B**), the X-axis indicates the log2 FC of the expression score in early PGCs (ePGCs) vs. postimplantation late epiblasts (postL-EPIs; primed pluripotent cells), while the Y-axis indicates that in preimplantation epiblasts (pre-EPIs; naïve pluripotent cells) vs. postL-EPIs. In (**B**), the macaque genes are colored red or orange if their orthologs in humans are present within 50 kb of active LTR5_Hs. *, *P* value < 1.0E-4; **, *P* value < 1.0E-15; NS, *P* value > 0.05. Human scRNA-Seq data [Messmer et al. (*27*) and Kojima et al. (*8*)] and macaque data [Sasaki et al. (*11*)] were used. (C and D) Enrichment of the human genes adjacent to LTR5_Hs (**C**) or their orthologs in macaques (**D**) among the genes upregulated in both PGCLCs/PGCs and naïve pluripotent cells. (E) Comparison of the genes upregulated in both PGCLCs/PGCs and naïve pluripotent cells between humans and macaques. The numbers in parentheses denote the numbers of genes adjacent to LTR5_Hs in the human genome or their orthologs in macaques. Only genes with ortholog information are included. The 95 genes (i) present in the vicinity of LTR5_Hs and (ii) that exhibited PGC- and naïve-specific expression patterns only in humans were defined as the genes likely to be regulated by LTR5_Hs. (F) Stratification of the genes that are likely to be regulated by LTR5_Hs according to the insertion date of the associated LTR5_Hs. On the various branches of the primate species tree, the numbers of the genes that are likely to be regulated by LTR5_Hs inserted in the corresponding branch are shown. The species tree was created with TimeTree (*64*). Nd, not determined. (G) Expression patterns of the genes likely to be regulated by LTR5_Hs. Genes related to glucose metabolism, genes related to oxidative phosphorylation, and genes whose proteins engage in PPIs with the proteins encoded by the genes above (see **Fig. S5B**) are annotated. Only genes exhibiting higher expression [mean expression (log2[CP10k+1] > 0.3] in both PGCLCs and ESCs are shown.

### Gradual progression of LTR5_Hs-mediated gene expression alterations during hominoid evolution

The LTR5_Hs insertions started after hominoid-OWM divergence and continued even after human-chimpanzee divergence (**Fig. S5A**) (*17*). This suggests that the gene expression alterations driven by LTR5_Hs have proceeded gradually during hominoid evolution. To address this possibility, we first determined the insertion dates of various LTR5_Hs loci (**Fig. S5A and Table S7**).

Subsequently, the genes that are likely to be regulated by LTR5_Hs (**Fig. 6E**) were classified according to the insertion dates of the associated LTR5_Hs loci (**Figs. S5A and 6F**). As shown in **Fig. 6F**, 24 out of 95 genes were associated with LTR5_Hs loci that were inserted in the common ancestor of the hominoid lineage (i.e., the branch “HCGOG” in **Fig. 6F**). On the other hand, the majority of the genes (63 genes) were associated with LTR5_Hs loci that were inserted after the common ancestor of Homininae (human, chimpanzee, and gorilla) (**Fig. 6F**). Of these, 34 genes were associated with human-specific LTR5_Hs loci (**Fig. 6F**). Finally, we examined the insertion dates of LTR5_Hs loci that are likely to regulate genes related to the glucose metabolism pathway (shown in **Figs. 5C and 5D**) and the genes encoding proteins that exhibit protein-protein interactions (PPIs) with the proteins encoded by the genes above (**Figs. S5B and 6G**). Most of the core glucose metabolic genes (4 out of 5 genes) were associated with the LTR5_Hs loci inserted in the common ancestors of Hominoidea or Hominidae (humans, chimpanzees, gorillas, and orangutans) (**Figs. S5B and 6G**). On the other hand, one of the core glucose metabolic genes (*ENO2*), the genes whose proteins have PPIs with the proteins of the core glucose metabolic genes above, and the genes related to oxidative phosphorylation (i.e., *NDUFAB1* and *NNT*) were associated with the LTR5_Hs inserted more recently (**Figs. S5B and 6G**).

LTR5_Hs insertions continued even after human speciation, and some LTR5_Hs loci are insertionally polymorphic in modern human populations (*41*). To address the roles of these polymorphic LTR5_Hs loci on the gene regulation in PGCs and naïve pluripotent cells, we identified LTR5_Hs loci that are present in the human reference genome (GRCh38) but not fixed in 2,504 human genomes used as a global reference of human genome variation (**Table S8**) (*42*). Subsequently, we checked whether these polymorphic LTR5_Hs loci overlap with the LTR5_Hs loci that are likely to regulate gene expression (**Fig. S6**). Of the 11 polymorphic LTR5_Hs loci detected, two are in the vicinity of genes (*FOLR1* and *TNK1*) upregulated in both PGCLCs and naïve ESCs. This suggests that very recent insertions of LTR5_Hs have also contributed to alterations of gene expression in these cell types. Together, these results support that the gene expression alterations driven by LTR5_Hs in PGCs and naïve pluripotent cells proceeded gradually during hominoid evolution.

## Discussion

Previous studies have suggested that there are similarities in gene expression between PGCs and naïve pluripotent cells. However, most of these studies have focused only on several key TFs and have not characterized the similarity at the whole-transcriptome level (*4, 5, 8-10, 21, 24, 26*). Furthermore, the regulatory basis underlying the gene expression similarity between these cell types has not been elucidated. In the present study, we characterized the transcriptome signature shared between PGCLCs and naïve ESCs in detail and illuminated the presence of a shared gene regulatory network between these cell types (**Fig. 1**).

We showed that numerous LTR5_Hs loci are activated as common enhancers in PGCLCs and naïve ESCs (**Figs. 3 and 5**). Although the enhancer activity of LTR5_Hs in naïve pluripotent cells has been reported in previous studies (*30, 31, 36, 37*), our data highlight the pleiotropic activity of the enhancers derived from LTR5_Hs, which likely contributes to the establishment of transcriptome similarity between PGCLCs and naïve ESCs. The results of our comparative transcriptome analysis between humans and macaques support the idea that LTR5_Hs insertions have altered the expression patterns of their adjacent genes in PGC- and naïve pluripotent cell-specific manners during hominoid evolution (**Fig. 6**). Furthermore, very recent insertions of LTR5_Hs loci (i.e., those which are human-specific or even polymorphic in the human population) likely also contribute to gene regulation in PGCs and naïve pluripotent cells (**Figs. 6F, 6G and S6**). Despite the centrality PGCLCs and naïve ESCs to maintenance of the germline (and by extension the species), our results suggest that gene expression in these cells may vary between humans based on polymorphisms in specific LTR5_Hs loci. Moreover, we found that LTR5_Hs loci are preferentially bound by key TFs shared between PGCLCs and naïve ESCs, such as *NANOG*, *TFAP2C*, *KLF4*, and *CBFA2T2*, suggesting that the enhancer activity of LTR5_Hs is likely regulated by these TFs (**Figs. 1C, 1F, 1H, and 4**) (*4, 5, 8, 9, 24, 26, 40*). These results further suggest that LTR5_Hs has incorporated its adjacent genes into the gene regulatory network driven by these TFs. Together, our data suggest that LTR5_Hs insertions gradually rewired the core gene regulatory network shared between PGCs and naïve pluripotent cells during hominoid evolution.

We found that genes related to the metabolism of carbohydrates, including glucose, were commonly upregulated in PGCLCs and naïve ESCs (**Fig. 1E**). In mice, the manner of glucose metabolism is similar between PGCs and naïve pluripotent cells (*28, 29, 43, 44*): mice PGCs and naïve pluripotent cells use both glycolysis and oxidative phosphorylation (i.e., both aerobic and anaerobic respiration, referred to as bivalent glucose metabolism), while primed pluripotent cells depend exclusively on glycolysis (i.e., anaerobic respiration). On the other hand, the manner of glucose metabolism in human PGCs is still unclear, although naïve human ESCs use bivalent glucose metabolism similar to that in naïve mouse ESCs (*28, 43, 45*). Together with the previous findings described above, our data suggest that human PGCLCs may also exhibit glucose metabolism similar to that of naïve ESCs (i.e., bivalent glucose metabolism), consistent with the case in mice. Notably, the manner of glucose metabolism affects the cellular identities of PGCs and naïve pluripotent cells in mice (*29*). Therefore, future functional studies seeking to characterize glucose metabolism in human PGCLCs are warranted.

Previous studies have demonstrated that naïve pluripotent cells in humans exhibit higher glycolytic activity than primed ones, while naïve pluripotent cells in mice and common marmosets (a New World Monkey) do not (*45, 46*). These findings suggest that glycolytic activity in naïve pluripotent cells was elevated in the hominoid or more ancestral lineages at least after the human-marmoset divergence. The data obtained in the present study suggest that the expression of genes related to glucose metabolism is likely controlled by LTR5_Hs in PGCs and naïve pluripotent cells and was likely upregulated in these cells during hominoid evolution (**Figs. 5C, 5D, and 6G**). Together, these findings raise the possibility that LTR5_Hs insertions are associated with the elevations in glycolytic activity in naïve pluripotent cells (and possibly in PGCs) during hominoid evolution. Since the manner of glucose metabolism substantially affects the identities of these cells (*29*), the enhancers derived from LTR5_Hs may affect the establishment or maintenance of these cells in humans by modulating glucose metabolism.

In conclusion, our data suggest that the core gene regulatory network shared between PGCs and naïve pluripotent cells has been finetuned by LTR5_Hs insertions during hominoid evolution. This gene regulatory network modification may contribute to the alterations in cellular characteristics, such as glucose metabolism, critical for the cellular identities of PGCs and naïve pluripotent cells. The present study provides insights into the germline evolution driven by selfish ERVs during hominoid evolution.

## Materials and Methods

### Bulk RNA-Seq of PGCLCs

The iPSC (9A13 XY) line used in this study was established in a previous study [Hwang et al. (*7*)]. iPSCs were cultured on plates coated with recombinant laminin-511 E8 (BG iMatrix-511 Silk, Peprotech, Cranbury, NJ) and were maintained under feeder-free conditions in StemFit Basic04 medium (Ajinomoto, Tokyo, Japan) containing basic FGF (Peprotech) at 37 °C under an atmosphere of 5% CO_2_ in air. For passaging or induction of differentiation, the cells were treated with a 1:1 mixture of TrypLE Select (Life Technologies, Waltham, MA) and 0.5 mM EDTA/PBS to enable their dissociation into single cells, and 10 mM ROCK inhibitor (Y-27632; Tocris, Abingdon, United Kingdom) was added.

PGCLCs were induced from iPSCs via iMeLCs as described previously [Sasaki et al. (*4*)] and purified using the surface markers EpCAM and INTEGRINα6. Total RNA was extracted from iPSCs and PGCLCs by using an RNeasy Micro Kit (Qiagen, Venlo, Netherlands) according to the manufacturer’s instructions. cDNA was synthesized using 1 ng of purified total RNA, and cDNA libraries were constructed for RNA sequencing by using a SMART-Seq HT Kit (Takara, Shiga, Japan) and a Nextera XT DNA Library Preparation Kit (Illumina, San Diego, CA) according to the manufacturers’ instructions. The libraries were sequenced using a single-end sequencing protocol on an Illumina NextSeq 500 instrument.

### Single-cell and bulk RNA-Seq analyses of human data

In the present study, read count matrices containing both human gene expression and subfamily-level TE expression data were prepared. To generate the count matrices, the human reference genome sequence (GRCh38/hg38) without ALT contigs was used. In addition, the gene and TE transcript annotation file (i.e., GTF file) generated in a previous study [Hwang et al. (*7*)] was used. Briefly, this annotation file contains the gene transcript annotations for GRCh38/hg38 from GENCODE version 22 (*47*) and the TE annotations for GRCh38/hg38 from the RepeatMasker output file (15-Jan-2014). TE loci with low reliability scores (Smith-Waterman scores < 2,500) were excluded. The annotation file is described in detail in and is available from the GitHub repository (https://github.com/TheSatoLab/TE_scRNA-Seq_analysis_Hwang_et_al/blob/master/CellRanger/input/hg38_TE_noAlt_uniq ue.gtf.gz).

Regarding the scRNA-Seq dataset for human early male germ cell development [Hwang et al. (*7*)], the read count matrix provided by Hwang et al. was used (https://github.com/TheSatoLab/TE_scRNA-Seq_analysis_Hwang_et_al/blob/master/count_matrix_data/vitro/data.merged.vitro.count.csv.gz). The read count matrix was generated using only reads that were uniquely mapped to the human reference genome.

A read count matrix was generated for the scRNA-Seq datasets for naïve and primed ESCs [Messmer et al. (*27*)] and for PGCLCs and iPSCs [Kojima et al. (*8*)]. The sequencing reads were downloaded and decrypted using the fastq-dump command in SRA Toolkit (https://ncbi.github.io/sra-tools/). If multiple FASTQ files were available for one single cell, the FASTQ files were concatenated. The sequencing reads were trimmed using Trimmomatic (version 0.39) (*48*) and subsequently mapped to the human reference genome using STAR (version 2.6.1c) (*49*) with the gene-TE transcript model described above. The read count matrix was constructed using featureCounts (version 1.6.3) (*50*). In this process, only reads that were uniquely mapped to the human reference genome were used.

Bulk RNA-Seq data for naïve and primed ESCs [Takashima et al. (*24*) and Theunissen et al. (*31*)] and for PGCLCs (original data obtained in the present study) were analyzed according to the same pipeline described in the above paragraph.

The read abundance of each TE subfamily was calculated by summing the read counts of TE loci belonging to the TE subfamily using an in-house Python script (https://github.com/TheSatoLab/TE_scRNA-Seq_analysis_Hwang_et_al/blob/master/make_count_matrix/script/sum_TE_count.subfamily.py). The counts per 10,000 (CP10k) value was calculated as the relative expression level, and the log2-transformed CP10k with a pseudocount of one (log2[CP10k+1]) value was subsequently computed.

Information on the RNA-Seq datasets analyzed in the present study is summarized in **Table S9**.

### Pseudotime analysis

Pseudotime analysis of the scRNA-Seq data for *in vitro*-derived human male germ cell development [Hwang et al. (*7*)] was performed using Monocle 2 (*34*) according to the procedures in the official tutorial (http://cole-trapnell-lab.github.io/monocle-release/docs/). The expression read count data were normalized under the negative binomial distribution assumption. In the pseudotime analysis, the 1,000 protein-coding genes that were the most differentially expressed during human male germ cell development were used. DDRTree was selected for the dimension reduction method.

### Data integration and dimension reduction analysis of scRNA-Seq data

Data integration between the Hwang et al. (*7*) and Messmer et al. (*27*) datasets followed by dimension reduction analysis was performed using Seurat 3 (version 3.2.2) (*51*) according to the scheme described in the Seurat tutorial (https://satijalab.org/seurat/vignettes.html). For each scRNA-Seq dataset, the expression data were normalized using SCTransform (*52*) by regressing out the total expression levels of mitochondrial genes. Subsequently, the datasets were integrated using the Seurat “anchoring” framework (*51*). In the data integration, the 3,000 most differentially expressed protein-coding genes in both datasets were used. The dimension reduction analysis was performed via uniform manifold approximation and projection (UMAP) (*51*) based on the integrated expression data. In the UMAP analysis, the first 30 principal components were used.

### Definition of the PGC-specific expression score

In this analysis, scRNA-Seq data for *in vitro*-derived human male germ cell development [Hwang et al. (*7*)] were used. The dataset includes data for a series of cells that were sequentially differentiated from iPSCs (iPSCs, iMeLCs, PGCLCs, MLCs, TCs, and T1LCs). As shown in the upper panel of **Fig. 1B**, the model representing the PGC-specific expression pattern was defined by a iPSC:iMeLC:PGCLC:MLC:TC:T1LC ratio of 0:0:1:0.5:0:0 (referred to as the model). In this model, the expression value of MLCs was set to 0.5 since it is known that the critical TFs of PGCs (e.g., *TFAP2A*, *TFAP2C*, *SOX17*, and *NANOG*) remain weakly expressed in MLCs (and in multiplying prospermatogonia cells, the *in vivo* counterparts of MLCs) (**Fig. 4D**) (*7*). As shown in the middle panel of **Fig. 1B**, for each gene and TE subfamily, the data representing the expression pattern were defined. Briefly, the relative expression (log2[CP10k+1]) values in the various cells were normalized as Z scores. Next, the mean expression values in the different cell types were calculated according to the Z scores above, and these mean expression values were rescaled to fit between 0 and 1. Here, a series of rescaled mean expression values is referred to as the data. Finally, as shown in the lower panel of **Fig. 1B**, the sum of squared residuals (SSR) between the model and the data was calculated, and the SSR value was subsequently −log10-transformed. This −log10(SSR) value was defined as the PGC-specific expression score. This analysis was performed using an in-house script (“calc_PGC_specific_expression_score.R”) available from the GitHub repository (https://github.com/TheSatoLab/LTR5_Hs_PGC_Naive_enhancer).

### Differential gene expression analysis

Differential gene expression analysis was performed using DESeq2 (version 1.26.0) (*53*). Only protein-coding genes were included in this analysis. Genes with relatively low expression levels (i.e., those with a 90^th^ percentile of reads per million value < 0.2) were excluded from the analysis. The statistical significance was calculated with the Wald test. The false discovery rate (FDR) value was calculated by the Benjamini-Hochberg (BH) method.

### Classification of protein-coding genes and TFs according to their expression patterns

In this analysis, the protein-coding genes that were expressed in the dataset of either Hwang et al. (*7*) or Messmer et al. (*27*) were used. Genes upregulated in PGCLCs were defined as the top 10% of genes with respect to the PGC-expression score among the genes expressed in the Hwang et al. dataset.

Genes upregulated in naïve ESCs were defined as the genes with log2 FC values > 1 and FDR values < 0.05 in the differential gene expression analysis between naïve ESCs vs. primed ESCs using DESeq2. According to the above definitions, the genes were classified as genes upregulated in both cell types, genes upregulated only in PGCLCs, genes upregulated only in naïve ESCs, and other genes.

The TFs shown in **Figs. 1C and 1F** were selected according to the following scheme. Briefly, a list of human TFs was downloaded from The Human Transcription Factors database (version 1.01; http://humantfs.ccbr.utoronto.ca/index.php) (*54*). *CBFA2T2* was manually added to the list of TFs. The listed TFs were classified as TFs upregulated in both cell types, TFs upregulated only in PGCLCs, TFs upregulated only in naïve ESCs, and other TFs according to the scheme described in the above paragraph. Of the TFs upregulated only in PGCLCs or only in naïve ESCs, the TFs with a mean log2(CP10k+1) value>0.6 in the corresponding cell type were selected. Of the TFs upregulated in both cell types, the TFs with a mean log2(CP10k+1) value >0.6 in either PGCLCs or naïve ESCs and with a mean log2(CP10k+1) value >0.3 in the other cell type were selected. In addition, *TFAP2A* was manually added to the list of the shown TFs. Information on the gene classification is summarized in **Table S1**.

### GO enrichment analysis

A gene-gene set association file including Molecular Signatures Database (MSigDB) canonical pathways and InterPro entries was used. The MSigDB canonical pathways were downloaded from MSigDB (http://software.broadinstitute.org/gsea/msigdb; version 6.1). InterPro entries were obtained from BioMart on the Ensembl website (www.ensembl.org; accessed on 13th February 2018).

The statistical significance values of the overlaps between the list of genes of interest and the predefined gene sets were calculated by one-tailed Fisher’s exact test. FDR values were calculated using BH method. As a universal (or background) set of genes, the protein-coding genes satisfying the following criteria were used: 1) genes included in the gene-gene set association file above and 2) genes whose expression was detected in either of the scRNA-Seq datasets [Hwang et al. (*7*) or Messmer et al. (*27*)].

In the GO enrichment analysis shown in **Fig. 1E**, the redundant gene sets whose members highly overlapped with each other were removed from the results. First, the gene sets with significant enrichment (FDR < 0.05) were ranked according to the odds ratio values. Second, if the gene members of a certain gene set highly overlapped with those of the upper-ranked gene sets, the gene set was removed from the results. Two gene sets were regarded as highly overlapping if the Jaccard index was greater than 0.5. This gene set filtering was performed with an in-house script (“rmRedundantGS_based_on_OR.py”) available from the GitHub repository (https://github.com/TheSatoLab/LTR5_Hs_PGC_Naive_enhancer).

### ATAC-Seq and ChIP-Seq analyses

Sequencing reads obtained from ATAC-Seq or ChIP-Seq were mapped to the human reference genome (GRCh38/hg38) using the BWA-MEM algorithm (version 0.7.17) (*55*). Reads mapped to the mitochondrial genome or with low mapping scores (mapping quality, MAPQ < 10) were removed using SAMtools (version 1.10) (*56*). In addition, PCR-duplicated reads were removed using Picard MarkDuplicates (version 2.18.16) (http://broadinstitute.github.io/picard/). Peak calling was performed using MACS2 callpeak (version 2.2.6) (https://pypi.org/project/MACS2/) with the threshold FDR < 0.05. For ChIP-Seq, the input control files were used in the peak calling step if the files were available. If >50,000 peaks were detected in one dataset, only the top 50,000 peaks with respect to statistical significance were used in the downstream analyses. Information on the analyzed data is summarized in **Table S9**.

### Identification of the open chromatin regions activated in PGCLCs or naïve ESCs compared to primed ESCs

First, the union (or merged) set of ATAC-Seq peaks between the two compared conditions (e.g., naïve ESCs vs. primed ESCs) was defined using the bedtools merge function (version v2.27.0) (*57*). Second, from the sequencing read alignment (BAM) file of each ATAC-Seq run, the reads that were assigned to the various merged peaks were counted using featureCounts (version 1.6.3) (*50*). Finally, the peaks (i.e., open chromatin regions) that were activated (log2 FC > 1; FDR < 0.05) in PGCLCs or naïve ESCs compared to primed ESCs were identified using DESeq2 (version 1.26.0) (*53*). Subsequently, the open chromatin regions were classified into those upregulated in both cell types, those upregulated only in PGCLCs, those upregulated only in naïve ESCs, and others.

### Genomic Regions Enrichment of Annotations Tool (GREAT) enrichment analysis

As shown in **Fig. 1G**, the enrichment of the open chromatin regions of interest (the open chromatin regions activated in both cell types, only PGCLCs, and only naïve ESCs) in the vicinity of the genes of interest (the genes upregulated in both cell types, only PGCLCs, and only naïve ESCs) was calculated according to the GREAT scheme (*58*). This method is explained in detail elsewhere (*59*). Briefly, regions of interest were defined as the regions within 50 kb of the transcription start sites (TSSs) of the genes of interest. Background regions were defined as the regions within 50 kb of the TSSs of all protein-coding genes. The lengths of the regions of interest and the background regions were calculated and referred to as Li and Lb, respectively. In the regions of interest and the background regions, the open chromatin regions were counted (referred to as counts of interest [Ci] and background counts [Cb], respectively). The fold enrichment value was calculated by dividing Ci/Cb by Li/Lb, and the statistical significance was evaluated using a binomial test. This analysis was performed using an in-house script (“great_pairwise.py”) available from the GitHub repository (https://github.com/TheSatoLab/LTR5_Hs_PGC_Naive_enhancer).

### Enrichment analysis of TF binding sites on the set of open chromatin regions of interest

A public ChIP-Seq dataset for 1,308 types of TFs provided by the GTRD (version 19.10) (*32*) was used. The ChIP-Seq peak data file “Homo sapiens_macs2_clusters.interval.gz” was downloaded from the database above (http://gtrd19-10.biouml.org/) on 20th May 2020. This file contains the single set of peaks (i.e., clustered peaks) for each TF. In this file, the peaks that had been computed for the same TF under the different experimental conditions (e.g., cell line, treatment, and study) were joined into clusters. For the various TFs, we detected overlaps between the TF binding sites and the open chromatin regions. Next, we classified the open chromatin regions according to (i) whether the open chromatin regions overlapped with the TF binding sites and (ii) whether the open chromatin regions belonged to a set of open chromatin regions of interest (i.e., those activated in both cell types, only PGCLCs, and only naïve ESCs). Subsequently, the odds ratios and *P* values were calculated with Fisher’s exact test. The FDR values were calculated with the BH method.

### Genomic permutation test

To calculate the fold enrichment of the overlaps between TE loci and a set of genomic regions of interest (e.g., ATAC-Seq peaks), randomization-based enrichment analysis (i.e., a genomic permutation test) was performed. The genomic regions of interest were randomized using the bedtools shuffle function (*57*); subsequently, the genomic regions of interest on TE loci in the randomized data were counted. This process was repeated 100 times, and the mean value of the counts in the randomized datasets was regarded as the random expectation value. The fold enrichment was calculated by dividing the observed count by the random expectation value. The P value was calculated according to the assumption of a Poisson distribution. The random expectation value was used as the lambda parameter of the Poisson distribution. This analysis was performed using an in-house script (calc_enrichment_randomized.great.py) available from the GitHub repository (https://github.com/TheSatoLab/LTR5_Hs_PGC_Naive_enhancer).

### Identification of the potential regulators of LTR5_Hs in PGCs and naïve pluripotent cells

We first identified the TFs that preferentially bind to LTR5_Hs using public ChIP-Seq data for 1,308 types of TFs provided by the GTRD (version 19.10) (*32*). For the various TFs, we calculated the fold enrichment of the TF-binding events on LTR5_Hs over the random expectation as well as the statistical significance using the genomic permutation test described in the above section. Next, we integrated the TF binding enrichment data with the expression pattern data of these TFs. To identify the potential regulators of LTR5_Hs in PGCs, the PGC-specific expression score defined in the above section was used. To identify the regulators in naïve pluripotent cells, the log2 FC values of the expression levels between naïve ESCs vs. primed ESCs computed using DESeq2 (*53*) were used. The potential regulators of LTR5_Hs were defined as the TFs satisfying the following criteria: (i) TFs that exhibited significant binding enrichment on LTR5_Hs (log2-fold enrichment > 2; FDR < 0.05; binding events > 20); (ii) for regulators in PGCLCs, TFs that were specifically upregulated in PGCLCs (in the top 10% with respect to the PGC-specific expression score; mean relative expression (log2[CP10k+1] > 0.4 in PGCLCs); and (iii) for regulators in naïve pluripotent cells, TFs that were specifically upregulated in naïve ESCs (log2[FC] > 2; FDR < 0.05; mean relative expression > 0.4 in naïve ESCs).

### Definition of the genes in the vicinity of active LTR5_Hs

The “active” LTR5_Hs loci, namely, the LTR5_Hs loci with transcriptomic or epigenetic signals, were defined. Specifically, LTR5_Hs loci with transcriptomic signals were defined as loci whose expression was detected in >0.5% of the cell population in any of the following scRNA-Seq datasets: (i) the PGCLC dataset of Hwang et al. (*7*), (ii) the PGCLC dataset of Kojima et al. (*8*), and (iii) the naïve ESC dataset of Messmer et al. (*27*). The LTR5_Hs loci with epigenetic signals were defined as loci that overlapped with the epigenetic signal peaks in any of the following ATAC-Seq or ChIP-Seq (targeting H3K27ac) datasets: (i) the PGCLC ATAC-Seq or ChIP-Seq dataset of Chen et al. (*21*) and (ii) the datasets of naïve ESCs in Pontis et al. (*30*). Information on the active LTR5_Hs loci is summarized in **Table S7**.

The genes in the vicinity of the active LTR5_Hs were also defined. The TSSs of the various transcripts for each protein-coding gene were extracted from the GENCODE gene annotation model (version 22) (*47*). The distance from the TSS of each gene to the closest LTR5_Hs copy was computed using the bedtools closest function (*57*). Subsequently, for each gene, the minimum distance from the TSS to the active LTR5_Hs copy was calculated. A gene in the vicinity of the active LTR5_Hs was defined as a gene within 50 kb of the minimum distance defined above.

### scRNA-Seq analysis of crab-eating macaque data and comparative transcriptome analysis between humans and macaques

For analysis of crab-eating macaque data, the reference genome (macFas5.fa), gene transcriptome annotation (genes/macFas5.ensGene.gtf; corresponding to the Ensembl 99 gene transcriptome annotation), and RepeatMasker output files (macFas5.fa.out) were downloaded from the University of California, Santa Cruz (UCSC) Genome Browser (http://hgdownload.soe.ucsc.edu/goldenPath/macFas5/bigZips/) on 23rd March 2020. The gene-TE transcript model for crab-eating macaques was constructed according to the same procedure used for humans. For the gene model, transcripts with the flag “retained intron” were excluded. For the TE model, TE loci with low reliability scores (i.e., Smith-Waterman scores < 2,500) were excluded. Additionally, the regions of TE loci overlapping with the gene transcripts were also excluded. The gene-TE transcript model was generated by concatenating the gene and TE models.

The scRNA-Seq dataset of early embryos and germ cells from crab-eating macaques [Sasaki et al. (*11*)] was analyzed. Briefly, the sequencing reads were trimmed using Trimmomatic (version 0.39) (*48*) and subsequently mapped to the reference genome using STAR (version 2.6.1c) (*49*) with the gene-TE transcript model above. The read count matrix was constructed using featureCounts (version 1.6.3) (*50*).

Gene ortholog information between humans and crab-eating macaques was downloaded from the Ensembl database (version 99) via BioMart (https://www.ensembl.org) on 23rd March 2020.

### Phylogenetic analysis of the LTR5 family

LTR5A, LTR5B, and LTR5_Hs loci with Smith-Waterman scores ≥ 2,500 were extracted from the RepeatMasker output file (15-Jan-2014; for GRCh38/hg38). Subsequently, the sequences of these LTR5 loci were extracted from the human reference genome (GRCh38/hg38) using the bedtools getfasta function (*57*). A multiple sequence alignment (MSA) of these LTR5 loci was constructed using MAFFT with the FFT-NS-i algorithm (version 7.407) (*60*). In the MSA, the alignment sites with <85% site coverage were eliminated using the in-house script “select_alignment_site.py” available from the GitHub repository (https://github.com/TheSatoLab/primate_A3_repertoire_and_evolution/blob/mai n/Trees/script). Subsequently, the sequences that had gaps in >15% of alignment sites were eliminated using the script above. In addition, tree-based filtering of the underlying dataset was performed prior to construction of a final tree. A preliminary tree was constructed, and phylogenetic outlier sequences, which have extremely long external branches (i.e., standardized external branch lengths > 3), were subsequently detected and discarded from the MSA used for final tree construction. The phylogenetic tree of LTR5 loci was reconstructed using RAxML (version 8.2.11) (*61*) with the GTRCAT model.

### Investigation of the distribution of orthologs of human LTR5 loci across Simiiformes

LiftOver chain files were downloaded from the UCSC Genome Browser (http://hgdownload.soe.ucsc.edu/goldenPath/hg38/liftOver/) (**Table S10**). Using the LiftOver program (http://genome.ucsc.edu/cgi-bin/hgLiftOver) and the LiftOver chain files, the genomic coordinates of LTR5 loci in the human reference genome were converted to those in another species with the option “Minmatch=0.5”. If the conversion was successful, we inferred that the orthologs of the LTR5 loci were likely present in the corresponding genome.

### Estimate of the insertion dates of LTR5_Hs loci and stratification of the genes likely to be regulated by LTR5_Hs according to the insertion dates

The insertion dates of the various LTR5_Hs loci were estimated according to information on both (i) the distributions of orthologous insertions across primates and (ii) the positions of LTR5 loci in the phylogenetic tree. Since there were a substantial number of missing values in the ortholog distribution information, we used phylogenetic information in addition to ortholog information to robustly estimate the LTR5_Hs insertion dates. First, LTR5_Hs loci were ordered according to the phylogenetic relationship (from older to younger). Second, using the framework of a sliding window analysis, the final positions of LTR5_Hs loci where more than three out of ten LTR5_Hs loci had orthologous insertions were determined for each primate of interest (chimpanzee, gorilla, orangutan, gibbon, macaque, and marmoset). For each species, LTR5_Hs loci that were older than the final LTR5_Hs copy were regarded as LTR5_Hs loci that were inserted before the divergence between humans and the corresponding species. Information on the estimated insertion dates is summarized in **Table S7**.

The genes that are likely to be regulated by LTR5_Hs were stratified according to the insertion dates of the associated LTR5_Hs loci. If the associated LTR5_Hs of one gene was not included in the phylogenetic tree of LTR5 loci, the gene was categorized as “not determined”. In addition, if multiple LTR5_Hs loci with distinct insertion dates were associated with one gene, the gene was also categorized as “not determined”.

### PPI network analysis

PPI network information for humans was downloaded from the Search Tool for the Retrieval of Interacting Genes/Proteins (STRING) database (version 11.0; “9606.protein.links.v11.0.txt.gz”) (*62*). The PPI links with confidence scores >400 were used for the analysis. The number of interacting partners of each gene was computed with the igraph package implemented in R (https://igraph.org/).

### Detection of LTR5_Hs insertions that are present in the human reference genome but not fixed in the human population

High-coverage whole genome sequencing (WGS) datasets in 1000 Genome Project (*42*) were downloaded from the following URL: ‘ftp://ftp.1000genomes.ebi.ac.uk/vol1/ftp/data_collections/1000G_2504_high_coverage/’. We searched WGS data for reads spanning the insertion site of of LTR5_Hs loci as follows. We first detected/annotated LTR5_Hs from GRCh38 using RepeatMasker with repeat sequence library provided from RepBase (version 24.01). We used the ‘-s -no_is’ options to sensitively detect LTR5_Hs. Next, we searched for reads skipping annotated LTR5_Hs, that is, reads mapping to the genomic regions flanking the LTR5_Hs insertion site (i.e. the predicted state/sequence of this locus prior to LTR5_Hs integration). We screened reads in the WGS datasets and extracted soft-clipped reads with ‘SA:Z’ tag. During this step, supplementary reads were excluded from analysis. We checked the mapped positions of the clipped and non-clipped regions on GRCh38. Here after, we refer to the clipped and non-clipped regions as to clipped_seq and non_clipped_seq, respectively. We next filtered out reads of which clipped_seq and non_clipped_seq are mapping to different chromosomes. Then we checked whether the clipped_seq and non_clipped_seq are mapping to flanking regions of an annotated LTR5_Hs locus. In this step, we considered that a read is a skipping read if both the clipped_seq and non_clipped_seq map to 25-nt from the ends of an annotated LTR5_Hs locus. We found 11 LTR5_Hs loci that are likely absent in at least one datasets. The mean count of skipping reads per LTR5_Hs locus in a single dataset ranged from 3.4 to 12.8. To exclude potential false positives due to any technical reasons, such as index hopping, we considered that an individual lacks at least one allele of a LTR5_Hs copy if two or more skipping reads were found at the LTR5_Hs locus.

### Data visualization

All data visualizations were performed in R (version 3.6.3). Heatmaps were drawn using ComplexHeatmap (*63*). The phylogenetic tree was visualized with ggtree (http://bioconductor.org/packages/release/bioc/html/ggtree.html). The PPI network was visualized using ggnet2 (https://briatte.github.io/ggnet/). The other data were visualized with ggplot2 (https://ggplot2.tidyverse.org/).

### Statistical analysis

Statistical analysis was performed in R (version 3.6.3). Statistical significance was evaluated by the two-tailed Wilcoxon rank sum test unless otherwise noted. FDR values were calculated by BH method.

## General

We would like to thank Mai Suganami (Institute of Medical Science, The University of Tokyo, Japan) for technical supports; Junna Kawasaki (Institute for Frontier Life and Medical Sciences, Kyoto University, Japan) for thoughtful comments. The super-computing resource, SHIROKANE, was provided by Human Genome Center, The Institute of Medical Science, The University of Tokyo, Japan. The results shown here are in part based on data generated by the 1000 Genome Project (https://www.internationalgenome.org/about/).

## Author contributions

J. Ito and K. Sasaki designed this study; J. Ito performed bioinformatics analyses; S. Kojima, K. Sato, and NF. Parrish supported bioinformatics analyses; Y. Seita performed experimental analyses; J. Ito prepared the figures; J. Ito wrote the initial draft of the manuscript; all authors contributed to data interpretation, designed the research, revised the paper, and approved the final manuscript.

## Data and materials availability

The RNA-seq data reported in this paper are available in GEO (https://www.ncbi.nlm.nih.gov/geo/query/acc.cgi?acc=GSE167570). The data produced in this study are available from the Mendeley Data Repository (http://dx.doi.org/10.17632/w5gfs9mdrr.1). The computational codes used in this study are available from the GitHub repository (https://github.com/TheSatoLab/LTR5_Hs_PGC_Naive_enhancer).

## Supplementary Figures

Fig. S1 TFs upregulated in both cell types, only PGCLCs, and only naïve ESCs

Fig. S2 Expression patterns of KZFPs

Fig. S3 Expression patterns of SVA transposons

Fig. S4 Pathway maps of glycolysis and glycogen breakdown

Fig. S5 Stratification of the genes likely to be regulated by LTR5_Hs according to the insertion date of the associated LTR5_Hs

Fig. S6 Potential roles of polymorphic LTR5_Hs insertions on the gene expression in PGCLCs and naïve ESCs

## Supplemental Tables

Table S1 Classification of protein-coding genes according to their expression patterns (related to **Fig. 1C**)

Table S2 GO enrichment analysis results for the three gene categories (genes upregulated in both cell types, genes upregulated only in PGCLCs, and genes upregulated only in naïve ESCs) (related to **Fig. 1E**)

Table S3 Identification of the potential regulators of LTR5_Hs in PGCLCs and naïve ESCs (related to **Fig 4**)

Table S4 Association of the expression patterns of genes and their distance from LTR5_Hs in the genome (related to **Fig. 5A**)

Table S5 Results of GO enrichment analysis using the genes that are present nearby LTR5_Hs and upregulated in both PGCLCs and naïve ESCs (related to **Fig. 5C**)

Table S6 Comparison of the genes upregulated in both PGCLCs/PGCs and naïve pluripotent cells between humans and macaques (related to **Fig. 6E**)

Table S7 Information on respective LTR5 loci (related to **Fig. S5A**)

Table S8 LTR5_Hs loci that are present in the human reference genome (GRCh38) but not fixed in the human population (related to **Fig. S6**)

Table S9 Sequencing dataset analyzed in the present study

Table Sx10 LiftOver chain files used in the present study

## Supplementary Figures

**Fig. S1.**
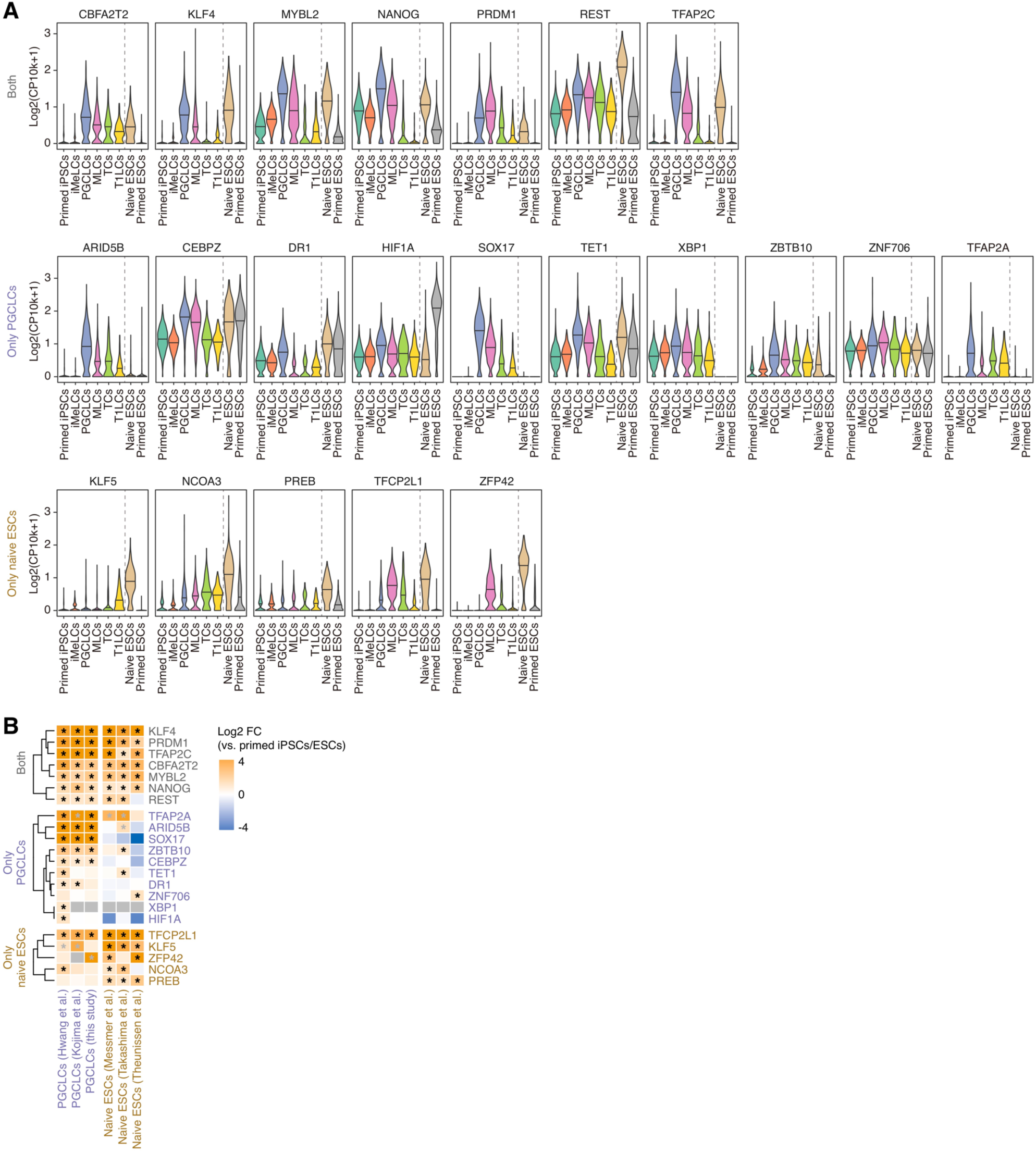
TFs upregulated in both cell types, only PGCLCs, and only naïve ESCs. (A) Expression levels in various cell types from scRNA-Seq data for male germline development [Hwang et al. (*7*)] and for naïve and primed ESCs [Messmer et al. (*27*)]. The results for the TFs annotated in Fig. 1C are shown. (B) Upregulation of TFs in PGCLCs and naïve ESCs observed across datasets. For the various datasets, the log2 FC values of the expression scores in PGCLCs vs. primed iPSCs or naïve ESCs vs. primed ESCs are shown. An asterisk denotes significant upregulation (FDR < 0.05; log2 FC > 1). A gray asterisk indicates that the expression level of the gene was not high (the mean expression level of the gene was below the 50th percentile for all expressed genes) even though significant upregulation was observed. For PGCLCs, the data of Hwang et al. (*7*) and Kojima et al. (*8*) were analyzed in addition to the original data in the present study. For naïve ESCs, the data of Messmer et al. (*27*), Takashima et al. (*24*), and Theunissen et al. (*31*) were analyzed.

**Fig. S2.**
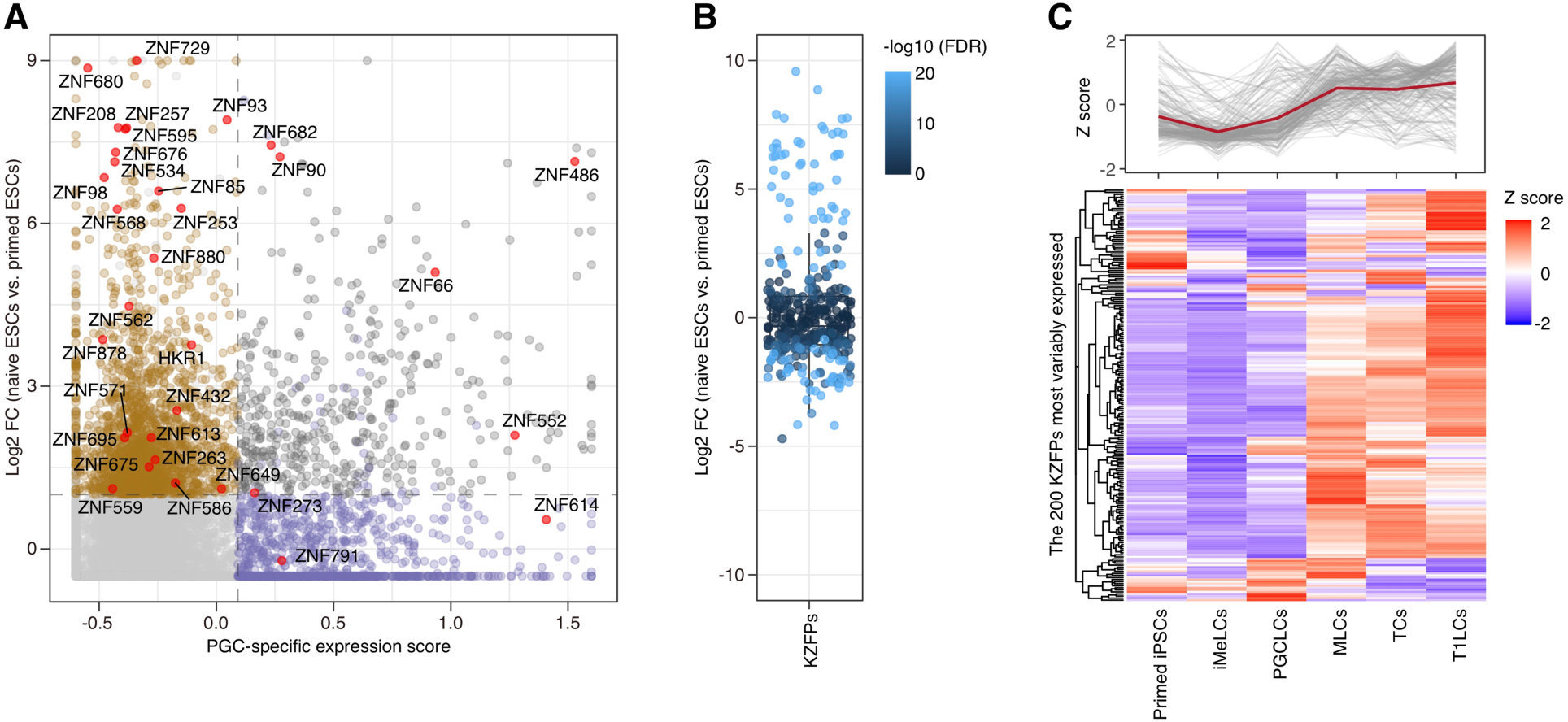
Expression patterns of KZFPs. (A) Classification of KZFPs according to their expression patterns. Highly expressed KZFPs in PGCLCs or naïve ESCs are annotated. The results for TFs other than KZFPs are shown in Fig. 1C. (B) Distributions of the log2 FC values of the expression scores of KZFPs in naïve ESCs vs. primed ESCs. The dot color denotes the statistical significance of the gene expression change. (C) Expression patterns of KZFPs during *in vitro*-derived human male germline development. The heatmap shows the relative mean expression values in the various cell types. The upper panel shows the transitions of the individual (gray) and mean (red) expression values.

**Fig. S3.**
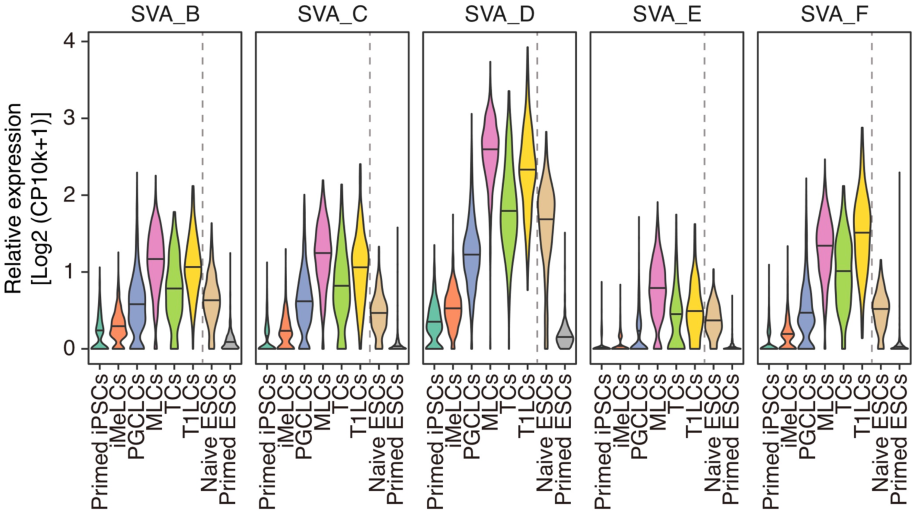
Expression patterns of SVA transposons. The results for the SVA transposons included in the heatmap in Fig. 2B are shown.

**Fig. S4.**
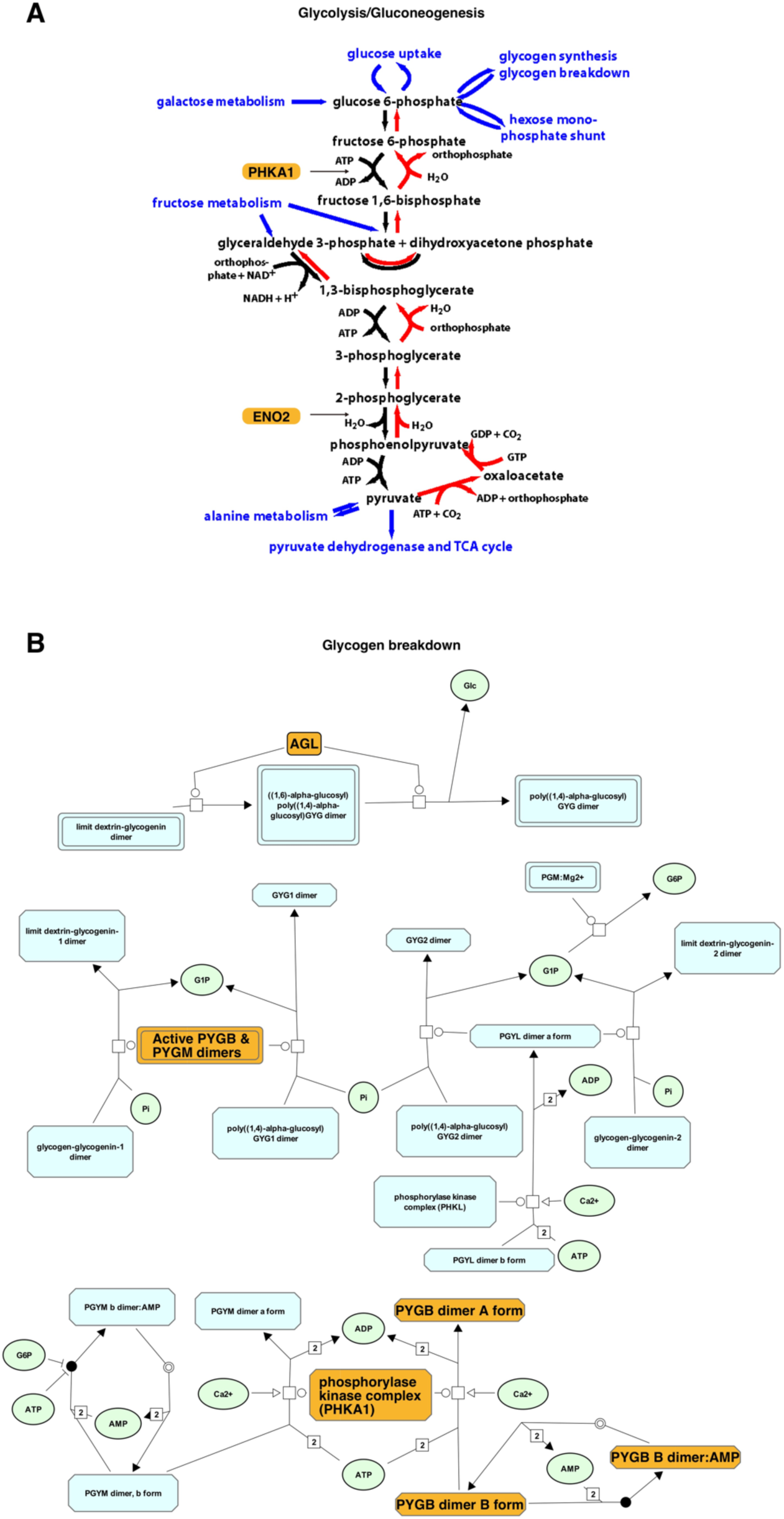
Pathway maps of glycolysis and glycogen breakdown. Pathway maps of glycolysis (**A**) and glycogen breakdown (**B**). Genes that are likely to be regulated by LTR5_Hs (i.e., *AGL*, *ENO2*, *PFKL*, *PHKA1*, and *PYGB*) are highlighted in orange. The pathway maps originated from the Reactome pathway database (https://reactome.org/) (*65*).

**Fig. S5.**
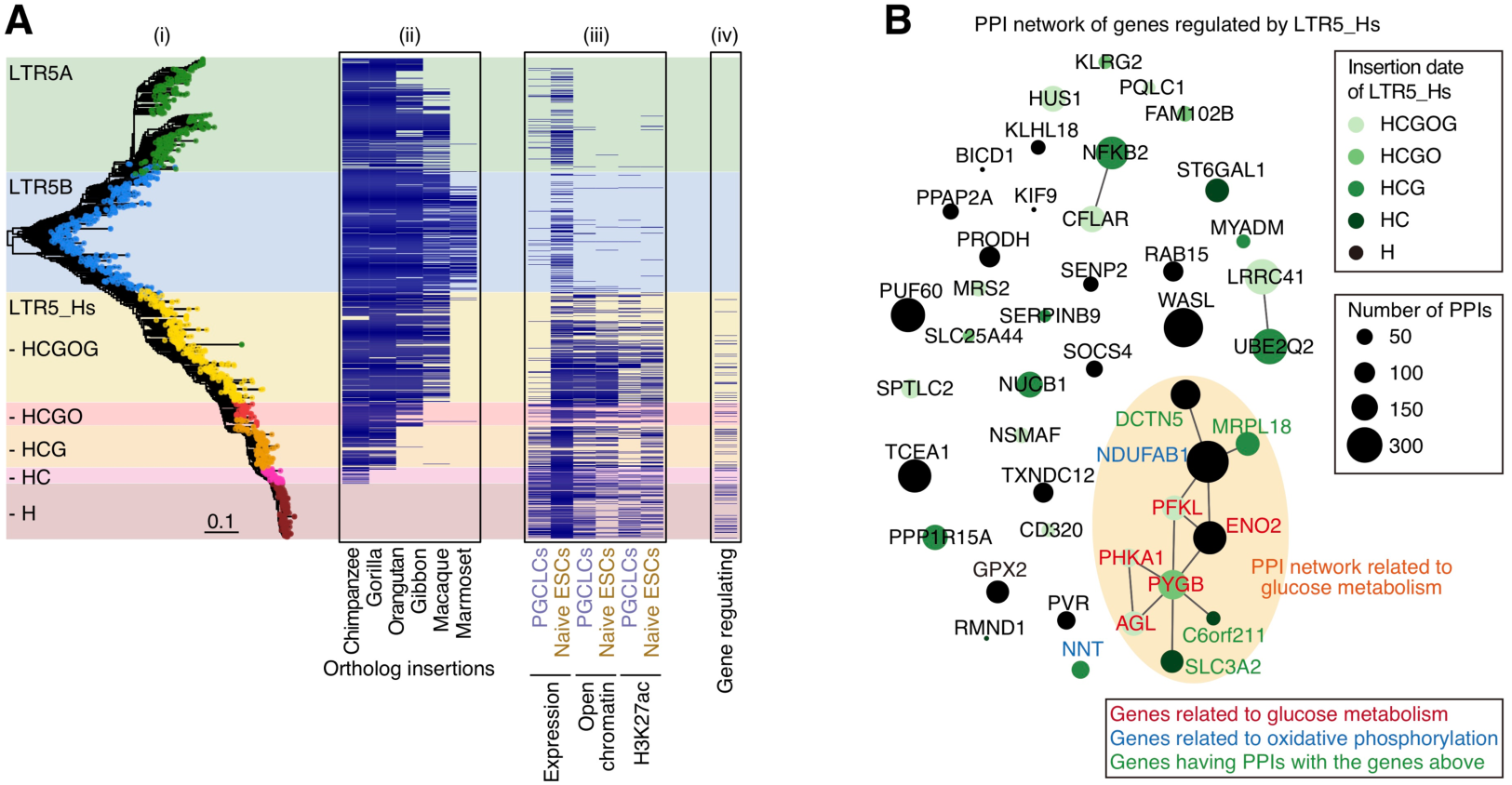
Stratification of the genes likely to be regulated by LTR5_Hs according to the insertion date of the associated LTR5_Hs. (A) Stratification of LTR5_Hs loci in the human genome according to their insertion dates. (i) Phylogenetic tree of the LTR5 family (including LTR5_Hs and related subfamilies [i.e., LTR5A and LTR5B]). (ii) Information on the distribution of orthologous insertions of LTR5 loci among primate genomes. According to the ortholog distribution and phylogeny, LTR5_Hs loci were stratified into five categories (HCGOG, HCGO, HCG, HC, and H). (iii) Epigenetic and transcriptomic statuses of various LTR5_Hs loci. (iv) LTR5_Hs loci that are likely to be associated with gene regulation. (B) PPI network for the genes likely to be regulated by LTR5_Hs. Only PPI links among the proteins encoded by the displayed genes are shown. The node color denotes the insertion date of the associated LTR5_Hs of the gene. The node size is proportional to the number of interacting partners in the whole PPI network. The glucose metabolism-related network is circled in orange. The PPI information originated from the STRING database (*62*).

**Fig. S6.**
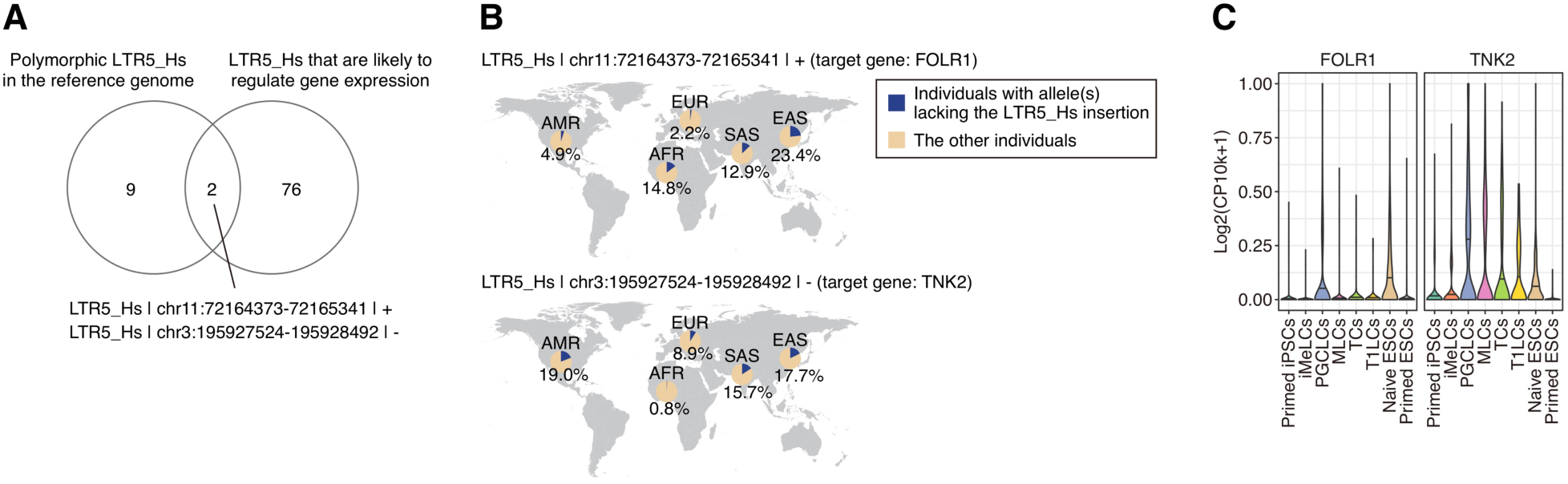
Potential roles of polymorphic LTR5_Hs insertions on the gene expression in PGCLCs and naïve ESCs. LTR5_Hs loci that are present in the human reference genome but not fixed in the human population (referred to as polymorphic LTR5_Hs loci) were identified using 1000 Genome Project datasets (*65*). Information on the polymorphic LTR5_Hs loci is summarized in **Table S8**. (A) Comparison of the polymorphic LTR5_Hs loci and the LTR5_Hs loci that are likely to regulate the gene expression in PGCLCs and naïve ESCs. The names of the overlapped LTR5_Hs loci are denoted (‘LTR5_Hs|chr11:72164373-72165341|+’ and ‘LTR5_Hs|chr3:195927524-195928492|-’). (B) Geographical prevalence of the polymorphic LTR5_Hs loci in respective human populations. Proportions of individuals with allele(s) lacking the LTR5_Hs insertion in respective populations are shown. AFR, African; AMR, Ad Mixed American; EAS, East Asian; EUR, European; SAS, South Asian. (C) Expression levels of the genes associated with polymorphic LTR5_Hs in various cell types.

